# A 3D brain unit model to further improve prediction of local drug distribution within the brain

**DOI:** 10.1101/688648

**Authors:** Esmée Vendel, Vivi Rottschäfer, Elizabeth C. M. de Lange

## Abstract

The development of drugs targeting the brain still faces a high failure rate. One of the reasons is a lack of quantitative understanding of the complex processes that govern the pharmacokinetics (PK) of a drug within the brain. While a number of models on drug distribution into and within the brain is available, none of these addresses the combination of factors that affect local drug concentrations in brain extracellular fluid (brain ECF).

Here, we develop a 3D brain unit model, which builds on our previous proof-of-concept 2D brain unit model, to understand the factors that govern local unbound and bound drug PK within the brain. The 3D brain unit is a cube, in which the brain capillaries surround the brain ECF. Drug concentration-time profiles are described in both a blood-plasma-domain and a brain-ECF-domain by a set of differential equations. The model includes descriptions of blood plasma PK, transport through the blood-brain barrier (BBB), by passive transport via paracellular and trancellular routes, and by active transport, and drug binding kinetics. The impact of all these factors on ultimate local brain ECF unbound and bound drug concentrations is assessed.

In this article we show that all the above mentioned factors affect brain ECF PK in an interdependent manner. This indicates that for a quantitative understanding of local drug concentrations within the brain ECF, interdependencies of all transport and binding processes should be understood. To that end, the 3D brain unit model is an excellent tool, and can be used to build a larger network of 3D brain units, in which the properties for each unit can be defined independently to reflect local differences in characteristics of the brain.

**Author summary:** Insights on how a drug distributes within the brain over both time and space are still limited. Here, we develop a ‘3D brain unit model’ in order to understand the factors that control drug concentrations within a small piece of brain tissue, the 3D brain unit. In one 3D brain unit, the brain capillaries, which are the smallest blood vessels of the brain, surround the brain extracellular fluid (ECF). The blood-brain barrier (BBB) is located between the brain capillaries and the brain ECF. The model describes the impact of brain capillary blood flow, transport across the BBB, diffusion, flow and drug binding on the distribution of a drug within the brain ECF. We distinguish between free (unbound) drug and drug that is bound to binding sites within the brain. We show that all of the above mentioned factors affect drug concentrations within brain ECF in an interdependent manner. The 3D brain unit model that we have developed is an excellent tool to increase our understanding of how local drug concentrations within the brain ECF are affected by brain transport and binding processes.

## 1 Introduction

The brain capillary bed is the major site of drug exchange between the blood and the brain. Blood flows from the general blood circulation into the brain capillary bed by a feeding arteriole and back by a draining venule. The rate at which drug molecules within the blood are exposed to the brain is determined by the brain capillary blood flow rate. Drug exchange between the blood plasma in the brain capillaries and the brain extracellular fluid (ECF) is controlled by the blood-brain barrier (BBB).

Drug distribution into and within the brain has been extensively summarized in a recent review [1]. In short, the BBB has great impact on the relationship between the concentration-time profiles of unbound drug in the blood plasma (blood plasma pharmacokinetics (PK)) and in the brain ECF (brain ECF PK). The BBB consists of brain endothelial cells that are held closely together by tight junctions. Unbound drug may cross the BBB by passive and/or active transport [2–10]. Passive transport is bidirectional and occurs by diffusion through the BBB endothelial cells (transcellular transport) and through the BBB tight junctions (paracellular transport). Active transport is unidirectional and can be directed inward (from the blood plasma to the brain ECF, active influx) or outward (from the brain ECF to the blood plasma, active efflux). Once having crossed the BBB, drug distributes within the brain ECF by diffusion. Diffusion within the brain ECF is hindered by the brain cells [11, 12]. This hindrance is described by the so-called tortuosity and leads to an effective diffusion that is smaller than normal (in a medium without obstacles). Moreover, a fluid flow, the brain ECF bulk flow, is present. The brain ECF bulk flow results from the generation of brain ECF by the BBB and drainage into the cerebrospinal fluid (CSF). Both diffusion and brain ECF bulk flow are important for the distribution of a drug to its target site, which is the site where a drug exerts its effect. In order to do induce an effect, a drug needs to bind to specific binding sites (targets). Only unbound drug, i.e. drug that is not bound to any components of the brain, can interact with its target [13, 14]. This is a dynamic process of association and dissociation, the so-called drug binding kinetics. These association and dissociation rates may affect the concentration of unbound drug at the target site [15, 16]. While the drug dissociation rate has been thought of as the most important determinant of the duration of interactions between a drug and its binding site [17], a more recent study shows that the drug association rate is equally important [16].

A number of models integrating several of the discussed processes of drug distribution into and within the brain is available, see for example [11, 12, 18–25] and [26]. The most recent and comprehensive brain drug distribution model is the physiologically-based pharmacokinetic model for the rat and for human [27, 28]. This model takes multiple compartments of the central nervous system (CNS) into account, including plasma PK, passive paracellular and transcellular BBB transport, active BBB transport, and distribution between the brain ECF, intracellular spaces, and multiple CSF sites, on the basis of CNS-specific and drug-specific parameters. However, it does not take into account distribution within brain tissue (brain ECF).

Here, we developed a 3D brain unit model, in which local brain drug distribution is explicitly taken into account. The 3D brain unit model encompasses blood plasma PK, the BBB, brain ECF, brain ECF bulk flow, diffusion, and binding to specific and non-specific binding sites within the brain. This 3D piece of brain tissue can be considered the smallest physiological unit of the brain in terms of drug transport. Within the 3D brain unit, drug is carried along with the blood plasma by the brain capillary blood flow and as such presented to the brain ECF. Drug distributes between the blood plasma and the brain ECF by transport across the BBB. Thereafter, drug distribution within the brain ECF is affected by diffusion, bulk flow and binding. We describe the distribution of drug within the brain ECF by a partial differential equation (PDE) and couple this to two ordinary differential equations (ODEs) to account for specific and non-specific drug binding.

The model builds on a proof-of-concept 2D brain unit model [29]. The 2D model is a basic model covering many essential aspects of drug distribution within the brain, including passive BBB transport, diffusion, brain ECF bulk flow, specific binding of a drug at its target site and non-specific binding of a drug to components of the brain. Here, brain cells are implicitly implemented by describing the hindrance the cells impose on the transport of a drug within the brain ECF in a tortuosity term. This model has enabled the study of the effect of drug properties and brain tissue characteristics on the distribution of a drug within the brain ECF and on its specific and non-specific binding behaviour of the drug.

The current *3D* brain unit model further improves the prediction of drug distribution within the brain. The third dimension improves the realistic features of the model as the brain is also 3D. Then, the brain capillary blood flow and active transport across the BBB, which are both important mechanisms of drug transport into the brain, are included. Here, we focus on one single brain unit. This allows for a thorough characterisation of drug distribution within one 3D brain unit before expanding to a larger scale.

In the remainder of this article, the mathematical representation of the characteristics of the 3D brain unit is introduced (section 2). There, we formulate the model (section 2.1) and the mathematical descriptions of the drug distribution within the blood plasma of the brain capillaries (section 2.2) and within the brain (section 2.3). In section 2.4 we formulate the model boundary conditions that describe drug exchange between the blood plasma and the brain ECF by passive and active BBB transport, as well as drug transport at the boundaries of the unit. In section 3, we study the effect of several factors on drug distribution within the brain ECF. In section 3.1, we evaluate the effect of the brain capillary blood flow velocity on local brain ECF PK in the 3D brain unit. Next, we evaluate the effect of active influx and efflux on local brain ECF PK (section 3.2). Then, in section 3.3 we show how the interplay between the brain capillary blood flow velocity, passive BBB permeability and active transport affects drug concentrations within the 3D brain unit. Finally, in section 4 we conclude our work and discuss future perspectives.

## 2 The 3D brain unit

The 3D brain unit represents the smallest piece of brain tissue that contains all physiological elements of the brain. The 3D brain unit is part of a larger network of 3D brain units, but here we focus on just one 3D brain unit that is fed by an arteriole and drained by a venule (Fig 1, left). The 3D brain unit is a cube in which the brain capillaries (represented by red rectangular boxes on the ribs) surround the brain ECF (Fig 1, left). The segments of red rectangular boxes protruding from the vertices from the 3D brain unit are parts of brain capillaries from neighbouring units. As such, each vertex connects three incoming brain capillaries to three outgoing brain capillaries, with the exception of the vertex connected to the arteriole and the vertex connected to the venule. These connect the arteriole to three outgoing brain capillaries and three incoming brain capillaries to the venule, respectively. A single 3D brain unit (Fig 1, middle) has a blood-plasma-domain (red) consisting of multiple sub-domains. These include the brain capillary domain where drug enters the unit (indicated by *U*_in_ in Fig 1), the domains representing the x-directed, y-directed and z-directed brain capillaries (indicated by *U*_x1*-*x4_, *U*_y1*-*y4_ and *U*_z1*-*z4_ in Fig 1) and the brain capillary domain where drug leaves the unit (indicated by *U*_out_ in Fig 1). Drug within the blood plasma is transported by the brain capillary blood flow. The brain capillary blood flow splits at the vertices of the unit, where brain capillary branching occurs (Fig 1, right).

**Fig 1.**
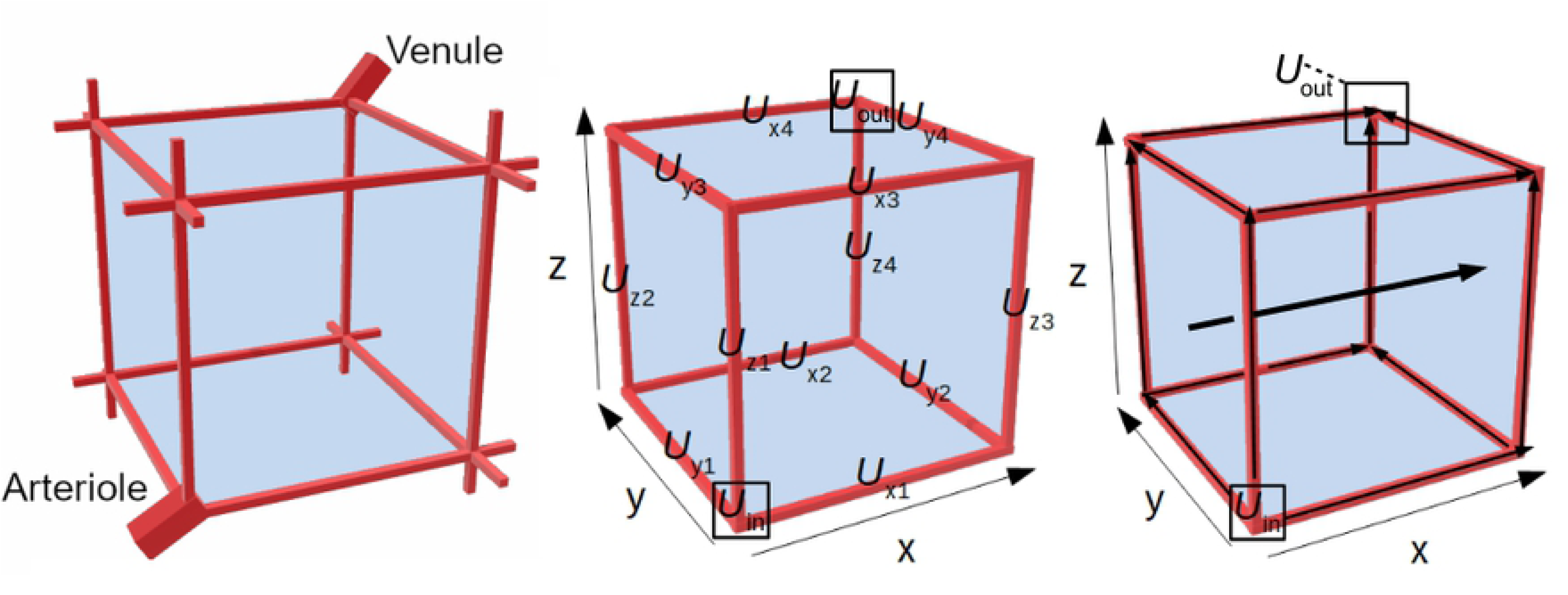
Sketch of the 3D model brain unit. Left: The structure represented by the 3D brain unit. An arteriole carries blood plasma (containing drug) into a brain capillary bed, that is connected to a venule that drains the blood plasma. The brain capillaries (red) surround the brain ECF (blue). Middle: the 3D brain unit and its sub-domains. The unit consists of a brain-ECF-domain (blue) and a blood-plasma-domain (red). The blood-plasma-domain is divided into several subdomains: *U*_in_ is the domain where the dose of absorbed drug enters the 3D brain unit, *U*_x1-x4_, *U*_y1-y4_ and *U*_z1-z4_ are the domains representing the x-directed, y-directed and z-directed capillaries, respectively. Right: Directions of transport in the model. The drug enters the brain capillaries in *U*_in_. From there, it is transported through the brain capillaries by the brain capillary blood flow in the direction indicated by the small arrows. Drug in the brain capillary blood plasma exchanges with the brain ECF by crossing the BBB. Drug within the brain ECF is, next to diffusion, transported along with brain ECF bulk flow (indicated by the bold arrow).

In developing the model, we make the following assumptions about drug distribution within the brain capillaries:

### Assumptions 1

i. The drug concentration within the blood plasma changes over time as a function of the rates of absorption (in case of oral administration) and elimination into and from the blood plasma.
ii. The blood carrying the drug flows into 3D brain unit by a feeding arteriole and leaves via a draining venule (Fig 1, left).
iii. The drugs enters the brain unit in the domain U_in_ (Fig 1, middle).
iv. The brain capillary blood flow is directed away from U_in_ (Fig 1, right).
v. Diffusion within the blood plasma is negligible compared to the brain capillary blood flow, hence drug is transported through the brain capillaries solely by the brain capillary blood flow.
vi. The brain capillaries are all equal in size and surface area. In addition, we assume that the volume of the incoming arteriole equals the volume of the three outgoing brain capillaries it connects to and that the volume of the outgoing venule equals the volume of the three incoming brain capillaries it connects to. Consequently, as the total volume of incoming blood vessels equals the total volume of outgoing blood vessels at each vertex (see Fig 1, left), the brain capillary blood flow velocity is by default equal in all brain capillaries.
vii. Drug within the blood plasma does not bind to blood plasma proteins. All drug within the blood plasma is in an unbound state and is able to cross the BBB.

Drug within the blood plasma of the brain capillaries crosses the BBB to exchange with the brain ECF. The BBB is located at the border between the brain capillaries (red) and the brain ECF (blue), see Fig 1. Drug exchange between the blood plasma and the brain ECF is described by passive and active transport across the BBB in both directions.

Within the brain ECF, we formulate:

### Assumptions 2

i. Drug within the brain ECF is transported by diffusion and brain ECF bulk flow.
ii. Cells are not explicitly considered, but only by taking the tortuosity (hindrance on diffusion imposed by the cells) into account.
iii. The brain ECF bulk flow is unidirectional. It is pointed in the x-direction, see the bold arrow in Fig 1 (right).
iv. All drug distributes within the brain ECF and we only have extracellular binding sites.
v. The total concentration of specific and non-specific binding sites is constant.
vi. The specific and non-specific binding sites are evenly distributed over the 3D brain unit and do not change position.
vii. The specific and non-specific binding sites lie on the outside of cells and the drug does not have to cross cell membranes in order to bind to binding sites.
viii. Drug binding is reversible and drugs associate and dissociate from their binding sites.

### 2.1 Formulation of the 3D brain unit

The 3D brain unit is a cubic domain, *U*, that represents a piece of brain tissue. We define U = {(x,y,z) ∈ ℝ^3^ | 0≤x≤ x_r_ ∧ 0≤y≤y_r_ ∧ 0≤z≤z_r_}. There, x_r_, y_r_ and z_r_ are constants that represent the length of one unit, which is then defined as *d*_cap_+2*r*, with *d*_cap_ the distance between the brain capillaries and *r* the brain capillary radius. In one brain unit, the brain capillaries, the BBB and the brain ECF are represented by the subsets *U*_pl_⊂*U, U*_BBB_⊂*U* and *U*_ECF_⊂*U*, respectively, such that *U* =*U*_pl_ ∪ *U*_BBB_ ∪ *U*_ECF_.

Within *U*_pl_, we define *U*_in_ as the domain where the blood plasma, containing drug, enters the 3D brain unit from a feeding arteriole. We define *U*_out_ as the domain where the blood plasma, containing drug, leaves the 3D brain unit to a draining venule. Additionally, we define the x-directed, y-directed and z-capillaries as the sets {*U*_xi_,i=1,..,4}, {*U*_yi_,i=1,..,4} and {*U*_zi_,i=1,..,4}. The brain capillaries are divided by the lines x=y (or y=z or x=z) and x+y=y_r_ (or y+z=z_r_ or x+z=z_r_), for which an example is shown in Fig 2. The only exceptions for this are the brain capillaries adjacent to *U*_in_ and *U*_out_, see below. The regions are defined are as follows:

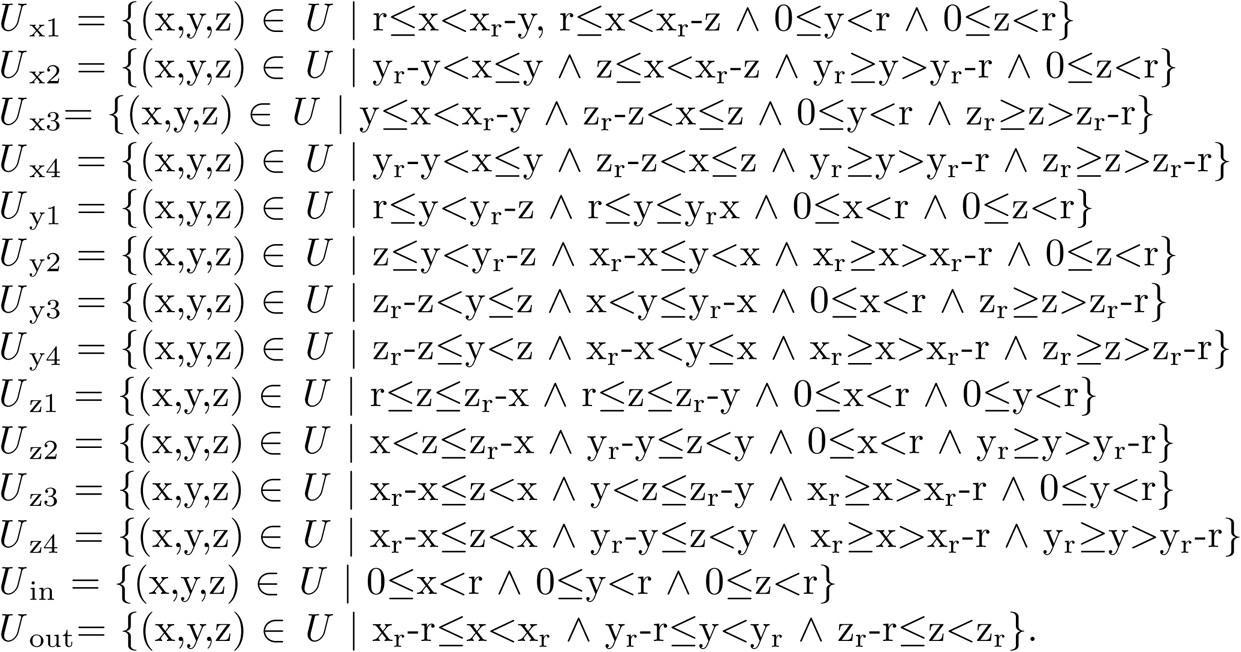

**Fig 2.**
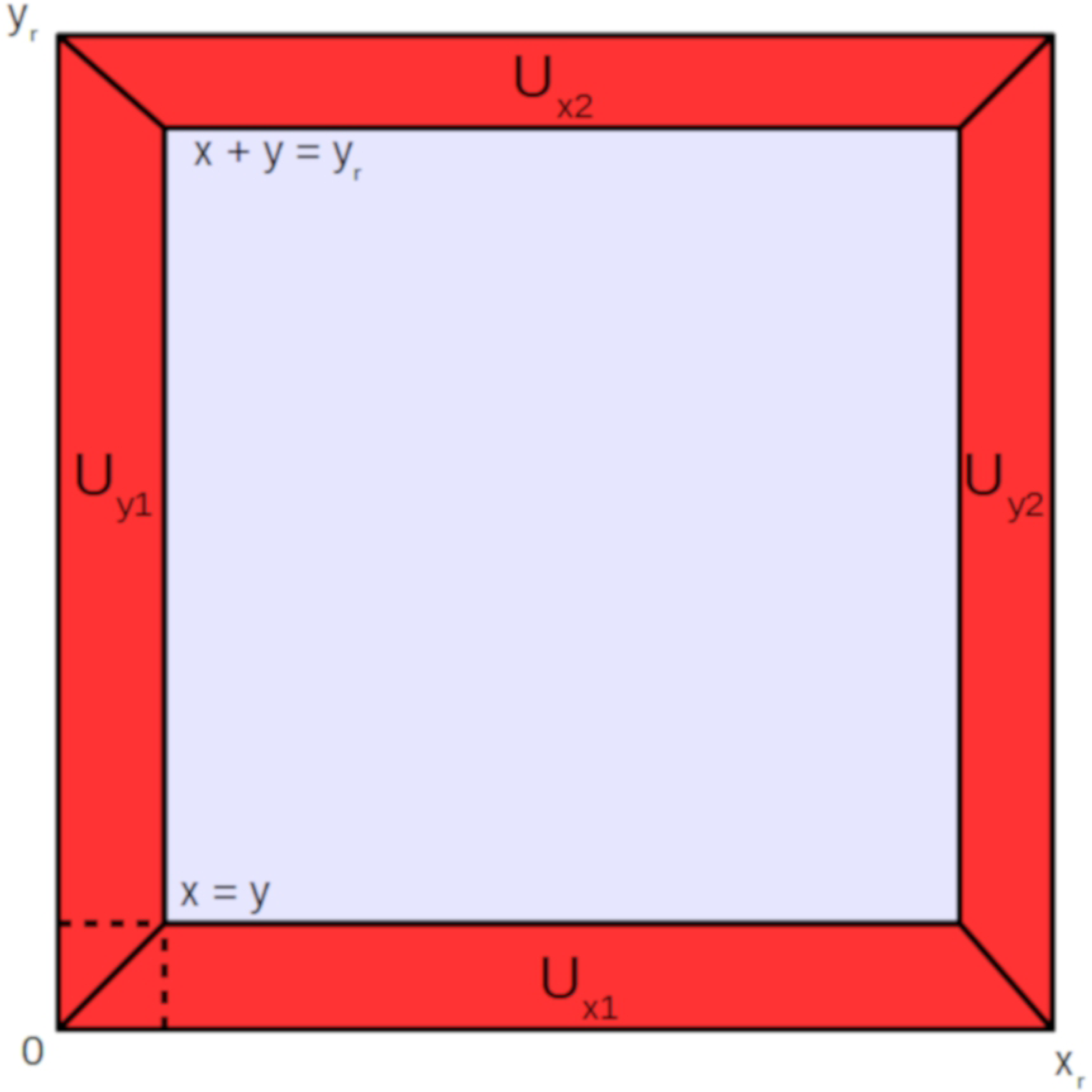
Front view of the 3D brain unit. Definitions of *U*_pl_ are given. The x-directed, y-directed and z-capillaries are divided by the lines x=y (or y=z or x=z) and x+y=y_r_ (or y+z=z_r_ or x+z=z_r_). The only exceptions for this are the brain capillaries adjacent to *U*_in_ and the brain capillaries adjacent to *U*_out_.

The BBB is represented by a subset *U*_BBB_⊂*U*, such that *U*_BBB_=*∂ U*_pl_ \ *∂ U*. This denotes the border between the blood plasma and the brain ECF, located at distance *r* from the edges of the 3D brain unit.

The brain ECF is represented by a subset *U*_ECF_⊂*U*, such that *U*_ECF_=*U*\(*U*_pl_∪*U*_BBB_). Within *U* we define the following quantities describing drug concentration:

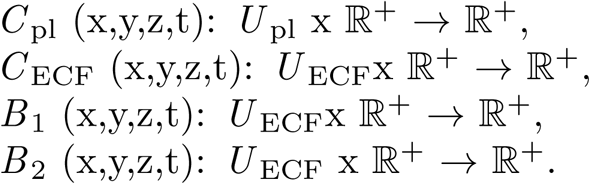

Here, *C*_pl_ is the concentration of unbound drug in the blood plasma, *C*_ECF_ is the concentration of unbound drug in the brain ECF, *B*_1_ is the concentration of drug in the brain ECF bound to specific binding sites and *B*_2_ is the concentration of drug in the brain ECF bound to non-specific binding sites.

### 2.2 Description of drug distribution in *U*_pl_

Based on assumption 1(i), we define the concentration of (unbound) drug within *U*_in_ by including parameters related to oral administration [30]:

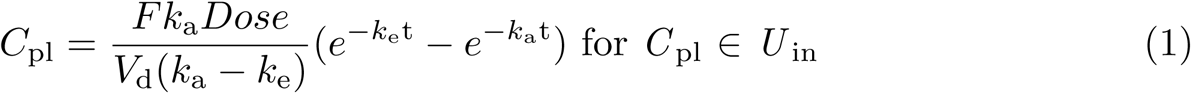

where *F* is the bioavailability of the drug, *k*_a_ the absorption rate constant of the drug, *k*_e_ the elimination rate constant of the drug, *Dose* the molar amount of orally administered drug, and *V*_d_ the distribution volume, which relates the total amount of drug in the body to the drug concentration in the blood plasma. We focus on oral administration but can also study other choices.

Additionally, based on assumptions 1(iv) and 1(v), we define:

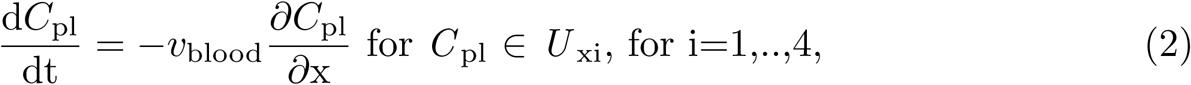

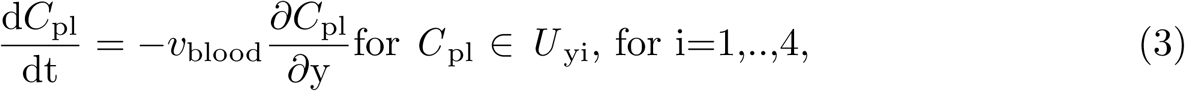

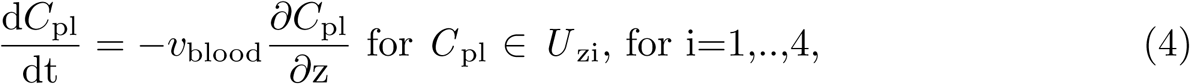

with *v* _blood_ the blood flow velocity within the brain capillaries and where the initial condition is given by

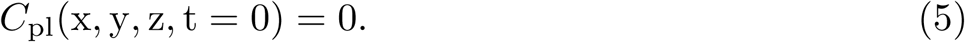

### 2.3 Description of drug distribution in *U*_ECF_

Based on assumptions 2, we describe the distribution of unbound and bound drug within *U*_ECF_ with the following system of equations:

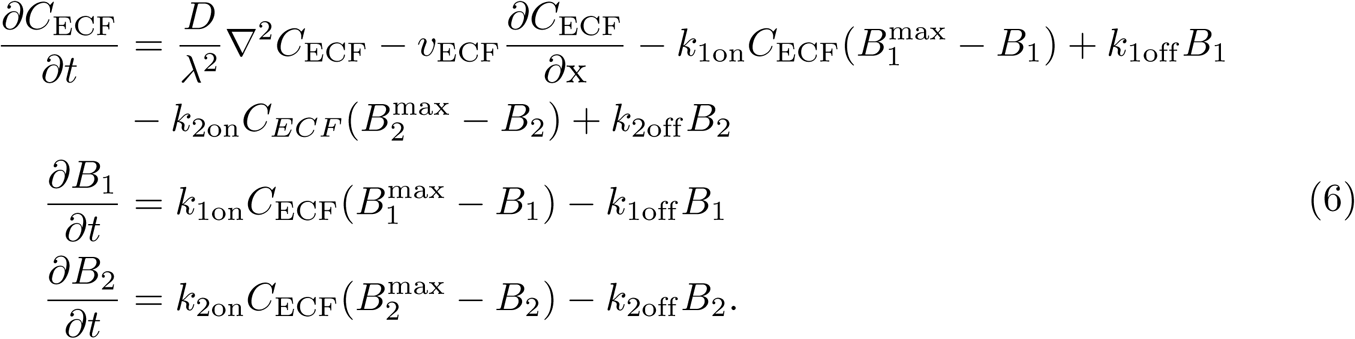

with initial conditions

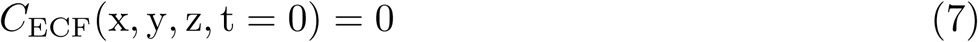

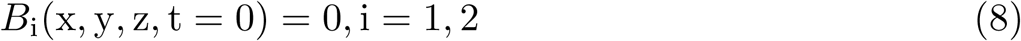

where *D* is the diffusion coefficient in a free medium, *λ* the tortuosity, *v* _ECF_ the (x-directed) brain ECF bulk flow, *B*_1_^max^,the total concentration of specific binding sites within the brain ECF, *k*_1on_ the association rate constant for specific binding, *k*_1off_ the dissociation rate constant for specific binding, *B*_2_^max^ the total concentration of non-specific binding sites within the brain ECF, *k*_2on_the association rate constant for non-specific binding and *k*_2off_ the dissociation rate constant for non-specific binding.

### 2.4 Boundary conditions

We formulate boundary conditions that describe the change in concentration of drug at the boundary between the blood-plasma-domain (*U*_ok_) and the brain-ECF-domain (*U*_ECF_), hence at *U*_BB_ as well as at the boundaries of the 3D brain unit (*U*_pl_∩*∂ U, U*_ECF_∩*∂U*).

#### 2.4.1 Drug exchange between *U*_pl_ and *U*_ECF_

We describe diffusive transport by the difference in drug concentrations in *C*_ECF_ and *C*_pl_, multiplied by the BBB permeability, *P*. In addition, we model active transport into and out of the brain ECF with Michaelis-Menten kinetics, similar to the approach of [6]. In total, this leads to:

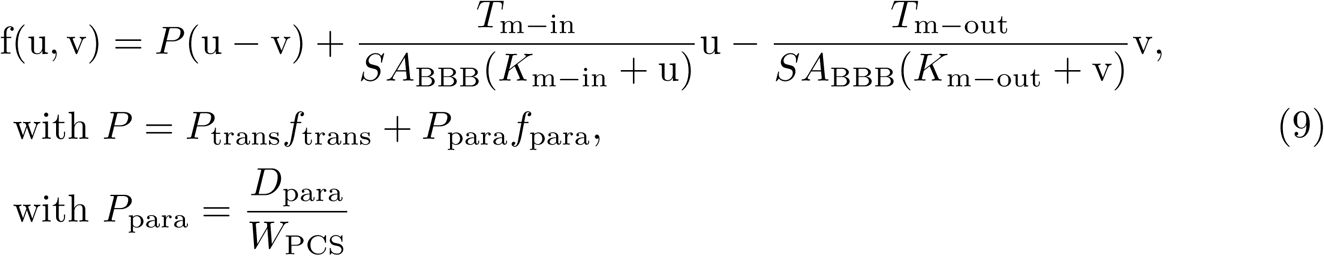

with u = *C*_pl_, v= *C*_ECF_, *P* _trans_ being the permeability through the brain endothelial cells, *f*_trans_ the fraction of the area occupied by the brain endothelial cells, *D*_para_the diffusivity of a drug across the paracellular space, *W*_PCS_ the width of the paracellular space, *f*_para_ the fraction of area occupied by the paracellular space, *T* _m-in_ the maximum rate of drug active influx, *T*_m-out_ the maximum rate of drug active efflux, *K*_m-in_ the concentration of drug at which half of *T*_m-in_ is reached, *K*_m-out_ the concentration of drug at which half of *T*_m-out_ is reached and *SA*_BBB_ the surface area of the BBB. Based hereon, we describe the loss or gain of unbound drug in the brain ECF due to BBB transport with the following boundary conditions (only those for the x direction are given, the ones for the y and z directions are similar):

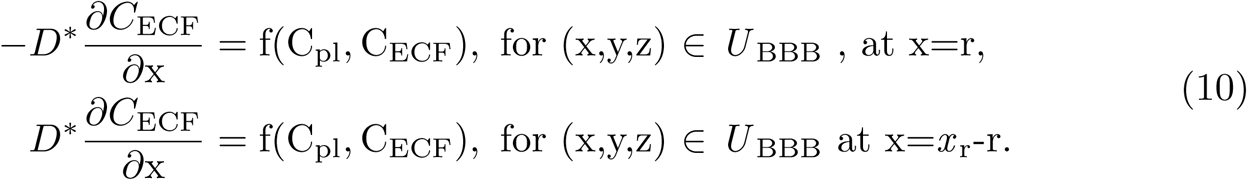

For the blood-plasma-domain, *U*_pl_, we use the reverse of (12) to describe drug transport across the BBB in the brain capillaries with the following boundary conditions:

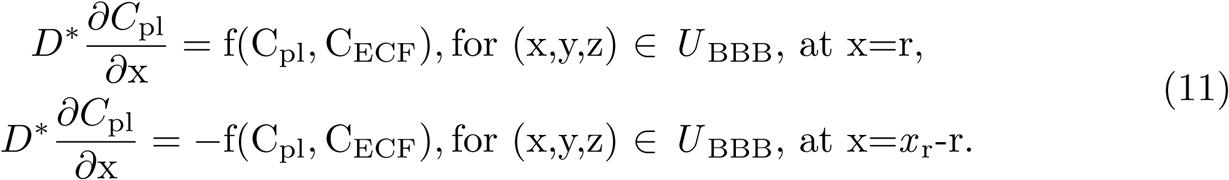

#### 2.4.2 Drug exchange at the faces of the 3D brain unit

We use additional boundary conditions to describe the drug concentrations at the sides of the domain. Since we assume that there is no diffusion in the blood plasma (see assumption 1(v)), we use the following boundary conditions:

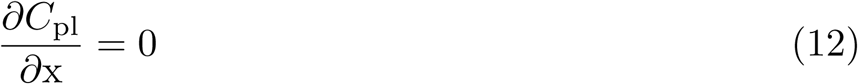

for (x,y,z) ∈ _pl_ *\ U*_out_ ∩*∂ U*, for x=0 and x=x_r_,

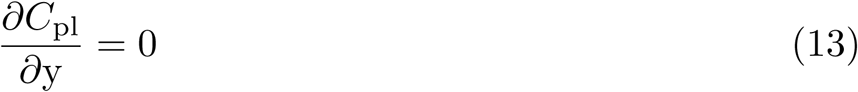

for (x,y,z) ∈ *U*_pl_ *\ U*_out_ ∩*∂ U*, for y=0 and y=y_r_,

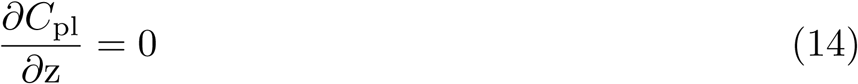

for (x,y,z) _pl_ \ *U*_out_ *∂ U*, for z=0 and z=z_r_.

In addition, we define:

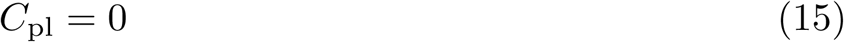

for (x,y,z) ∈ *U*_out_ ∩*∂ U*.

We formulate the condition at the boundaries of the 3D brain unit as follows:

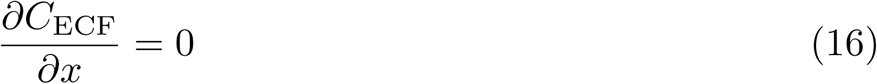

for *U*_ECF_∩*∂ U*.

### 2.5 Model parameter values and units

The dimensions of the 3D brain unit are based on the properties of the rat brain. The model is suitable for data from human or other species as well, but we have chosen for the rat as for this species most data is available. The distance between the brain capillaries in the rat brain is on average 50 *µ*m, while the brain capillaries have a radius of about 2.5 *µ*m [31–34]. Therefore, we set the radius of the brain capillaries, *r*, to 2.5 *µ*m and the dimensions of the 3D brain unit in the x, y and z directions, *x* _r_, *y* _r_ and *z* _r_ respectively, to 55 *µ*m.

In our model, we use Eq (2)–(6) to describe drug concentration within the blood plasma, with boundary conditions described in Eq (13)-(17). We describe the concentration of drug within the brain ECF with Eq (7)–(9) with boundary conditions described in (11),(12) and (18). The range of values we use for the parameters in the model as well as their units are given in Table 1 below. This range is based on values found in the literature (from experimental studies), which we also give in the table. The literature does not provide values on the kinetic parameters related to non-specific binding kinetics (*B*_2_^max^, *k*_2on_ and *k*_2off_). Therefore, we base the choices of these values on earlier articles that assume that drug binding to specific binding sites is stronger than to non-specific binding sites, while non-specific binding sites are more abundant [29, 35, 36].

**Table 1.**
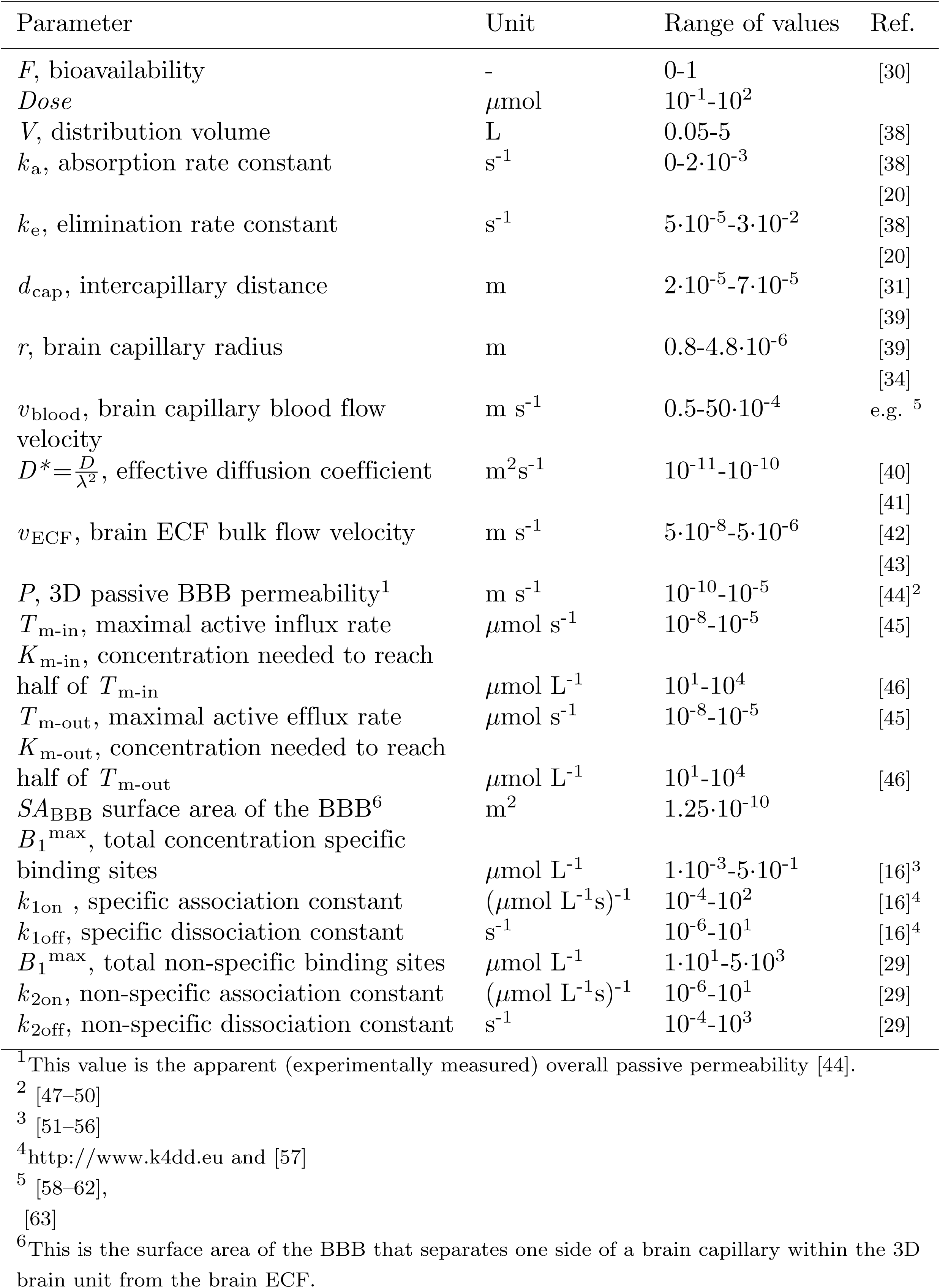
3D brain unit model parameters and their units, for rat brain. The physiological range of values of the parameters is given. These are based on references from the literature.

## 3 Model results

We study the distribution of a drug within the 3D brain unit by plotting its concentration-time profiles within the brain ECF (brain ECF PK). In addition, we study the distribution of the drug within the 3D brain unit. We first nondimensionalise the system of equations and boundary conditions by scaling all variables by a characteristic scale, see S1 Appendix for details. Next, in order to perform simulations, we discretise the nondimensionalised system spatially, using a well-established numerical procedure based on finite element approximations [37]. We present the results using the parameters with dimensions. The output of the simulations are the concentrations of free, specifically bound and non-specifically bound drug, given in *µ*mol L^−1^ over time (s). The model can easily be used to study a specific drug by choosing the parameter values that are specific for this drug, provided that parameter values for this drug are known. In the present study, however, we choose to study generic parameter values that are in the middle of the physiological ranges given in Table 1. This allows us to perform a sensitivity analysis and study the effect of parameter values at both extremes of the physiological range on the behaviour of the model. We use, unless otherwise indicated, the parameter values that are given in Table 2. In the following sections, we show the impact of the brain capillary blood flow velocity (*v*_blood_) in the absence of active transport (section 3.1), the impact of active transport (section 3.2) and the impact of *v*_blood_ and active transport combined (section 3.3) on blood plasma and brain ECF PK and brain ECF drug distribution. We give the concentration-time profiles of unbound drug, specifically bound drug and non-specifically bound drug in the middle of *U*_ECF_, where 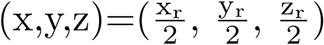 as well as those of unbound drug in the blood plasma in the middle of *U*_x1_, where 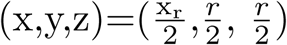, on a log-scale versus time. Drug distribution profiles are given for cross-sections of the entire (x,y,z)-domain of the 3D brain unit for various times.

**Table 2.** 3D brain unit model default parameter values and their units. The values are for a hypothetical drug and are all within the physiological ranges given in Table 1.

| Parameter | Unit | Value |
| --- | --- | --- |
| $F$ | - | 1 |
| $Dose$ | $\mu\text{mol}$ | 0.5 |
| $k_a$ | $\text{s}^{-1}$ | $2 \cdot 10^{-4}$ |
| $k_e$ | $\text{s}^{-1}$ | $5 \cdot 10^{-5}$ |
| $V$ | L | 0.2 |
| $d_{\text{cap}}$ | m | $5 \cdot 10^{-5}$ |
| $r$ | m | $2.5 \cdot 10^{-6}$ |
| $v_{\text{blood}}$ | $\text{m s}^{-1}$ | $5 \cdot 10^{-4}$ |
| $D^*$ | $\text{m}^2 \text{s}^{-1}$ | $0.5 \cdot 10^{-10}$ |
| $v_{\text{ECF}}$ | $\text{m s}^{-1}$ | $0.5 \cdot 10^{-6}$ |
| $P$ | $\text{m s}^{-1}$ | $0.1 \cdot 10^{-7}$ |
| $T_{\text{m-in}}$ | $\mu\text{mol s}^{-1}$ | $0 \cdot 10^{-7}$ |
| $T_{\text{m-out}}$ | $\mu\text{mol s}^{-1}$ | $0 \cdot 10^{-7}$ |
| $K_{\text{m-in}}$ | $\mu\text{mol L}^{-1}$ | $1 \cdot 10^2$ |
| $K_{\text{m-out}}$ | $\mu\text{mol L}^{-1}$ | $1 \cdot 10^2$ |
| $SA_{\text{BBB}}$ | $\text{m}^2$ | $1 \cdot 10^{-10}$ |
| $B_1^{\text{max}}$ | $\mu\text{mol L}^{-1}$ | $5 \cdot 10^{-2}$ |
| $k_{1\text{on}}$ | $(\mu\text{mol L}^{-1} \text{s})^{-1}$ | 1 |
| $k_{1\text{off}}$ | $\text{s}^{-1}$ | $1 \cdot 10^{-2}$ |
| $B_1^{\text{max}}$ | $\mu\text{mol L}^{-1}$ | $5 \cdot 10^1$ |
| $k_{2\text{on}}$ | $(\mu\text{mol L}^{-1} \text{s})^{-1}$ | $1 \cdot 10^{-2}$ |
| $k_{2\text{off}}$ | $\text{s}^{-1}$ | 1 |

### 3.1 The effect of the brain capillary blood flow velocity on brain ECF PK within the 3D brain unit

The impact of the brain capillary blood flow velocity, *v*_blood_, on brain ECF PK within the 3D brain unit is evaluated. Parameters are as in Table 2 and we thus assume that there is no active transport, i.e. *T*_m-in_=0 and *T*_m-out_=0. Here, we focus on the effect of *v*_blood_ on brain ECF PK in the middle of the 3D brain unit. We show the concentration-time profiles of unbound, specifically bound and non-specifically bound drug (*C*_ECF_, *B*_1_ and *B*_2_, respectively) within the 3D brain unit on a larger time-scale, for several values of *v*_blood_. We do so for the default value of the passive permeability *P* (*P* =0.1·10^−7^ m s^−1^), in Fig 3 (left), as well as for a high value of *P* (*P* =100·10^−7^ m s^−1^), in Fig 3 (right). The lowest value of *v*_blood_ is outside the known physiological ranges (see Table 1), but we choose it as *v*_blood_ is predicted to mostly impact drug concentrations in the brain when *P* is much higher than *v*_blood_ [64, 65]. The total passive permeability, *P*, includes both transcellular and paracellular permeability. The paracellular space may increase due to disruption of the tight junctions in certain disease conditions, thereby allowing larger molecules to pass through and increasing paracellular transport [66, 67]. We can tune our model and separate between transcellular and paracellular transport, as we do in S2 Appendix. In the current section we proceed with the total passive BBB permeability.

**Fig 3.**
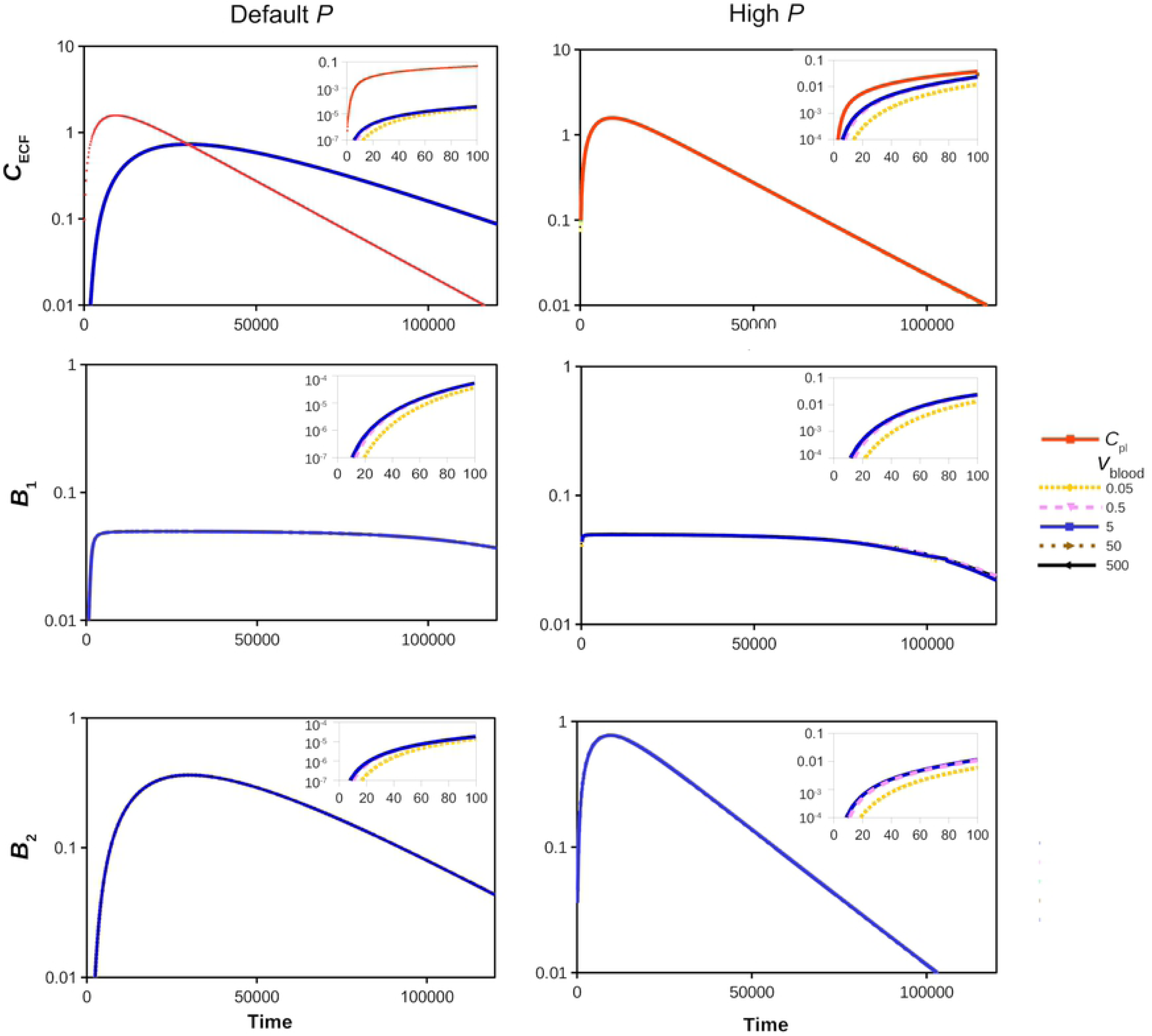
The effect of the brain capillary blood flow velocity, *v*_blood_ (m s^−1^), on the log PK of *C*_pl_ (red) and *C*_ECF_ (top), *B*_1_ (middle) and *B*_2_ (bottom) for a default (*P* =0.1·10^−7^m s^−1^) (left) and a high (*P* =100·10^−7^m s^−1^) (right) value of *P*. Values of *v*_blood_ are set at 0.05*·*10^−4^ m s^−1^, 0.5·10^−4^ m s^−1^, 5·10^−4^ m s^−1^, 50·10^−4^ m s^−1^ and 500·10^−4^ m s^−1^, as is depicted by different colours, where drug concentrations for the default value of *v*_blood_ (*v*_blood_=5·10^−4^ m s^−1^) are shown in blue. All other parameters are as in Table 2. The insets in each sub-figure show the PK for a shorter time.

Fig 3 shows that *v*_blood_ does not impact long-time behaviour of *C*_ECF_, *B*_1_ and *B*_2_. The insets in Fig 3 demonstrate that *v*_blood_ impacts short-time (t=0-100 s) behaviour only when it has extremely low values (*v*_blood_*≤*0.5*·*10^−4^ m s^−1^), as depicted in the insets of Fig 3 by the yellow and purple lines, respectively. The impact of *v*_blood_ on *C*_ECF_, *B*_1_ and *B*_2_ is independent of the values of *P* (compare the left and right insets of Fig 3). The effects of *P* on drug concentrations within the brain ECF are similar to those found with our proof-of-concept 2D model [29]: for a high value of *P*, the attained values of *C*_ECF_ and *B*_2_ are higher and follow *C*_pl_, while their decay is faster than for a low value of *P*. In addition, the *≥*90% maximum value of *B*_1_, i.e. values of *B*_1_ that are more than ≥ 90% of the maximum value attained during the simulation (*B*_1_ ≥90% max(*B*_1_)), is attained shorter for a high value of *P* than for a low value of *P*.

From the results shown in Fig 3 we conclude that the effects of *v*_blood_ on brain ECF PK are minimal. According to the Renkin-Crone equation [64, 65], the brain capillary blood flow affects drug *influx*, depending on the permeability of the BBB. This is also demonstrated by our model, and we show that *v*_blood_ affects drug influx across the BBB in S3 Appendix.

The plots in Fig 4a show the changes in concentration of drug within the blood plasma over a short time-range (t=5 to t=25). There, *C*_pl_ is plotted along the capillaries starting at *U*_in_ (where drug enters the unit) to *U*_out_ (where drug exits the unit). We measure the distance from *U*_in_, where the total distance between these points is 150 *µ*m. Drug can be transported along several pathways, but in Fig 4a the values of *C*_pl_ are given along the pathway indicated in Fig 4b. When *v*_blood_=0.5 (left), there are clear differences between *C*_pl_ in *U*_in_ (Distance=0) and *C*_pl_ in the opposite corner (Distance=150) at the time-points shown. However, as *C*_pl_ increases over time, the differences in *C*_pl_ become small relative to the value of *C*_pl_. Fig 4c shows the distribution profiles of unbound drug within the 3D brain unit at t=5 for different values of *v*_blood_. There, darker shades of red and blue correspond to higher concentrations of unbound drug in the blood plasma and the brain ECF, respectively. When *v*_blood_=0.5*·*10^−4^ m s^−1^, the transport time of drug between *U*_in_ and the opposite corner is higher than when *v*_blood_=5*·*10^−4^ m s^−1^. This is depicted in Fig 4c, where at t=5, drug concentrations within *U*_pl_ are equal for a high brain capillary blood flow velocity (*v*_blood_=50*·*10^−4^ m s^−1^), while local differences in *C*_pl_ still exist for a low value of *v*_blood_ (*v*_blood_=0.5*·*10^−4^ m s^−1^). The value of *v*_blood_ also affects local concentrations of *C*_ECF_. For a low value of *v*_blood_ (*v*_blood_=0.5*·*10^−4^ m s^−1^), values of *C*_ECF_ at t=5 are overall low, but highest in the corners closest to *U*_in_. For higher values of *v*_blood_ (*v*_blood_=5*·*10^−4^ m s^−1^ and *v*_blood_=50*·*10^−4^ m s^−1^), *C*_ECF_ at t=5 is overall higher, but again highest in the corner close to *U*_in_.

**Fig 4.**
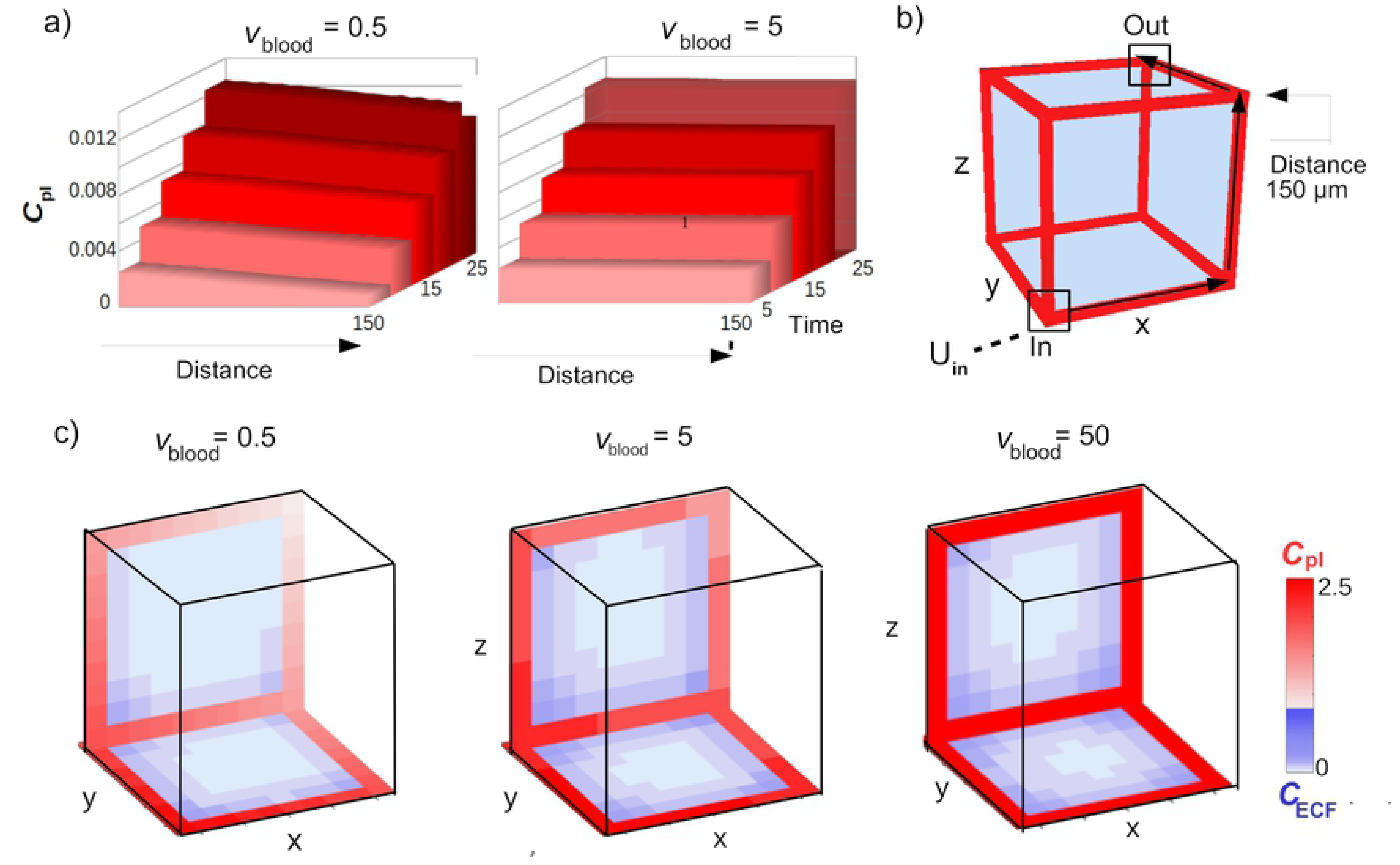
Changes in *C*_pl_ and *C*_ECF_ due to the effect of *v*_blood_. While *v*_blood_ is varied from 0.05·10^−4^ m s^−1^ to 50·10^−4^ m s^−1^, all other parameter values are as in Table 2. a) The pathway from *U*_in_ to *U*_out_ along which *C*_pl_ is plotted. b) *C*_pl_ is plotted against time (timepoints from 5 to 25) along the distance shown in (a). c) Distribution profiles of *C*_pl_ (red) and *C*_ECF_ (blue) of the 3D brain unit at t=5. Darker shades of red and blue correspond to higher values of *C*_pl_ and *C*ECF, respectively.

### 3.2 The effect of active transport on the drug concentrations within the brain ECF

Active transport kinetics are regulated by the maximal transport rate (*T*_m_) and the concentration of drug needed to reach half of the maximal transport rate (*K*_m_), see section 2.4.1. We first focus on active influx, such that *T*_m-out_=0. We vary *T*_m-in_, which denotes the maximal rate of active transporters moving drug from the blood plasma *into* the brain ECF. Fig 5 shows the effects of increasing values of *T*_m-in_ (starting at *T*_m-in_=0, i.e. no active influx) on *C*_ECF_ (top), *B*_1_ (middle) and *B*_2_ (bottom). Fig 5 (top) reveals that an increased value of *T*_m-in_ correlates with increased concentrations of *C*_ECF_. The time to the peak of *C*_ECF_ is not affected by the value of *T*_m-in_. Fig 5 (middle) shows that *T*_m-in_ does affect the time during which the specific binding sites are saturated. We find that 90% max(*B*_1_) is attained longer for a higher *T*_m-in_. Fig 5 (bottom) shows that higher values of *T*_m-in_ correlate with higher values of *B*_2_ and thus a greater occupancy of non-specific binding sites. The non-specific binding sites within the brain ECF become saturated with drug when *T*_m-in_ is sufficiently high (*T*_m-in_=100*·*10^−7^ *µ*mol s^−1^). To evaluate the effect of active efflux on drug concentrations within the brain ECF, we repeat our simulations with *T*_m_ directed outward, i.e. with *T*_m-out_=0-100·10^−7^ *µ*mol s^−1^ and *T*_m-in_=0. Fig 6 (top) shows that *C*_ECF_ decreases faster for higher values of *T*_m-out_, corresponding to more active efflux. Fig 6 (middle) reveals that *T*_m-out_ affects the time during which specific binding sites are saturated: the time at which *B*_1_ attains 90% max(*B*_1_) is smaller for a high value of *T*_m-out_. For sufficiently high values of *T*_m-out_, the binding sites do not become saturated. Fig 6 (bottom) shows that *B*_2_ is similarly affected by active efflux as *C*_ECF_.

**Fig 5.**
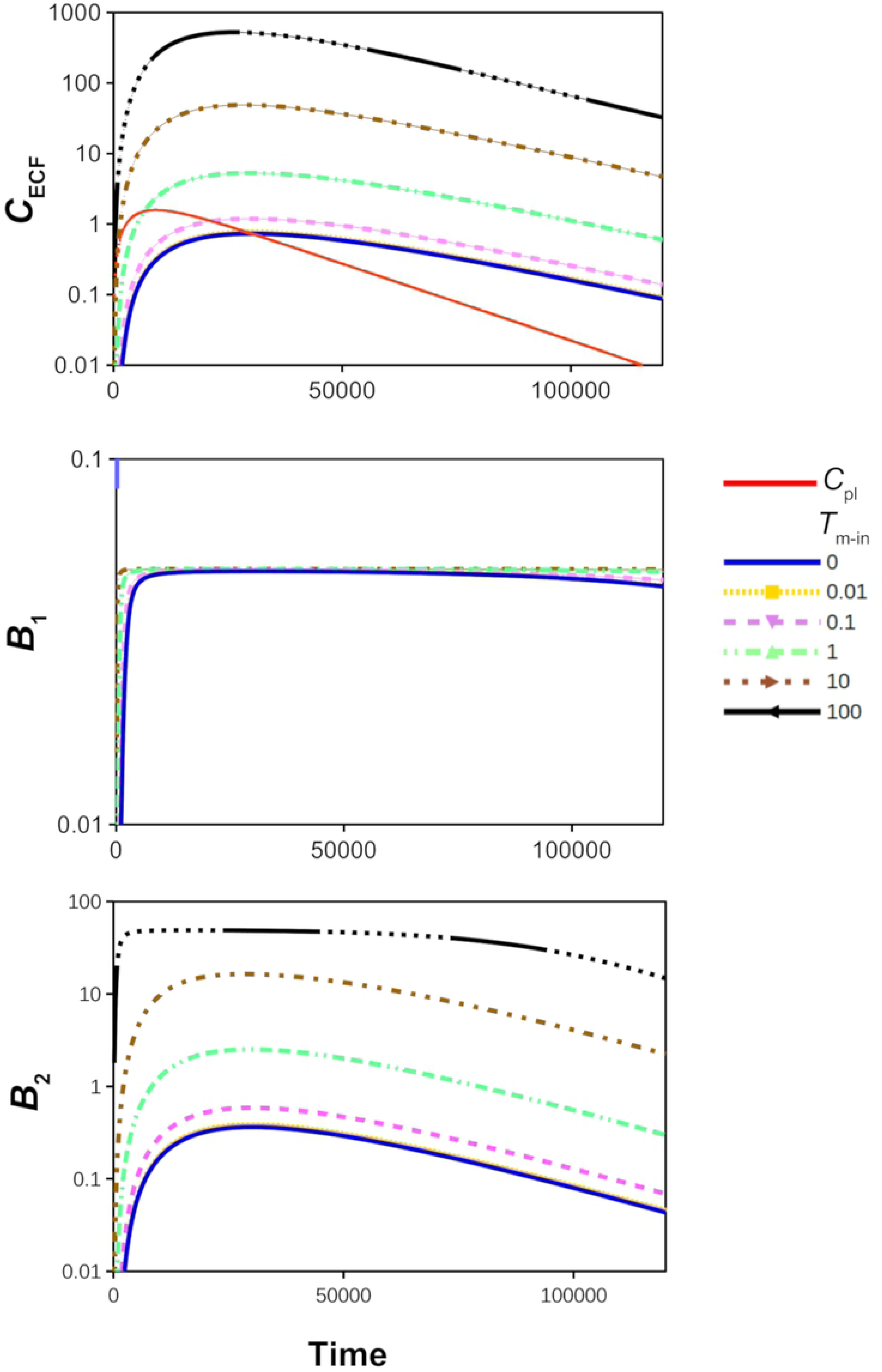
The effect of active influx on the log concentration-time profiles of drug in the brain ECF, relative to those in the blood plasma. Top: unbound drug in the brain ECF (*C*_ECF_) compared to unbound drug in the blood plasma (*C*_pl_, red curve). Middle: drug bound to its target sites (*B*_1_). Bottom: drug bound to non-specific binding sites (*B*_2_). The value of *T*_m-in_ is changed from 0 to 100*·*10^−7^ *µ*mol s^−1^. The rest of the parameters are as in Table 2.

**Fig 6.**
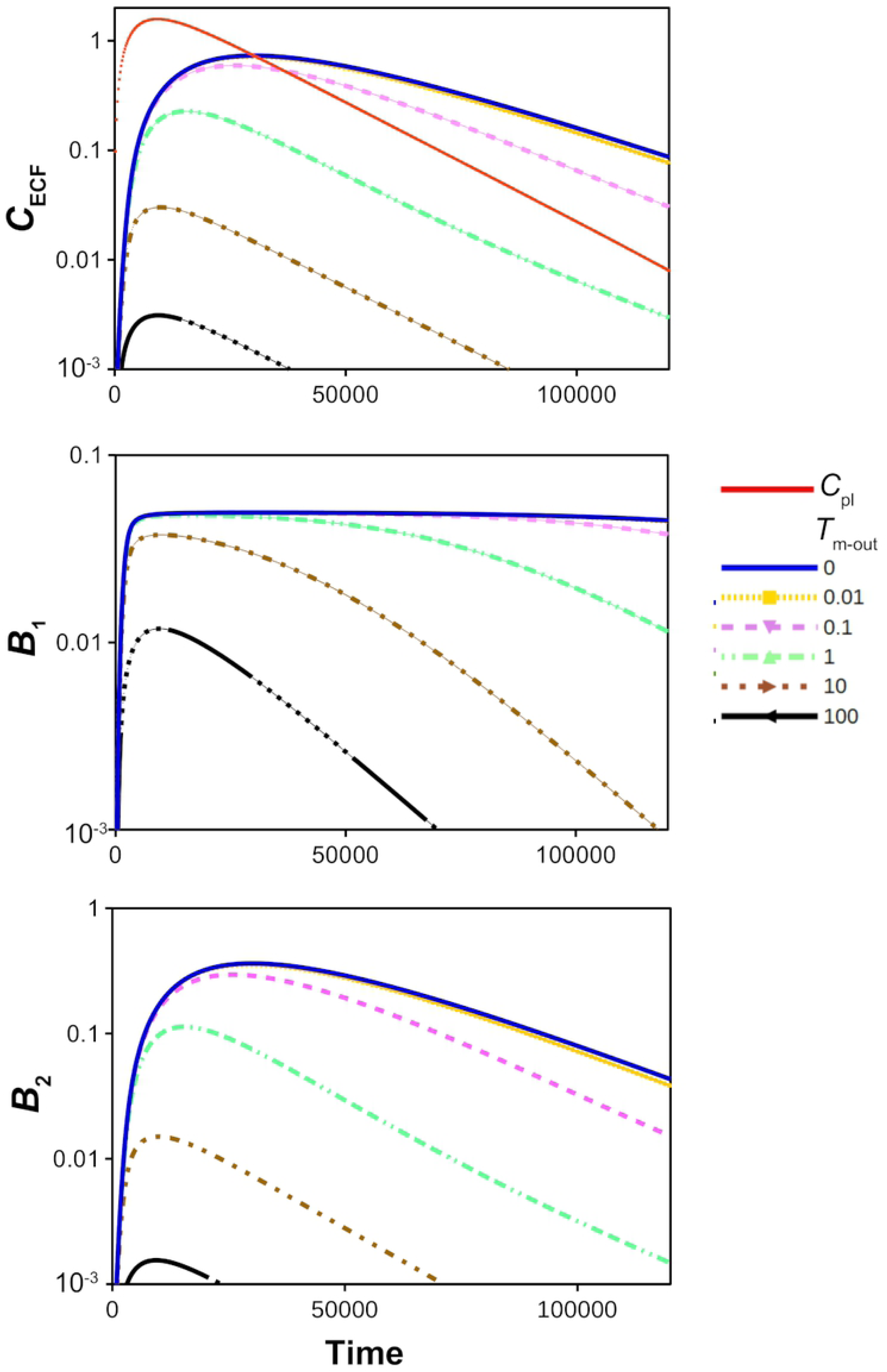
The effect of active efflux on the log concentration-time profiles of drug in the brain ECF, relative to those in the blood plasma. Top: unbound drug in the brain ECF (*C*_ECF_) and unbound drug in the blood plasma (*C*_pl_, red curve). Middle: drug bound to its target sites (*B*_1_). Bottom: drug bound to non-specific binding sites (*B*_2_). The value of *T*_m-out_ is changed from 0 to 100*·*10^−7^ *µ*mol s^−1^. The rest of the parameters are as in Table 2.

### 3.3 The effect of the brain capillary blood flow velocity in the presence of active transport

In section 3.1 we have shown that both the passive BBB permeability, *P*, and the brain capillary blood flow velocity, *v*_blood_, affect dug brain ECF PK in the absence of active transport. Here, we study how *P* and *v*_blood_ combined with active transport affect drug PK within the brain ECF. Fig 7 shows the log plot of *C*_ECF_ for *v*_blood_=5*·*10^−4^ m s^−1^ (top) and *v*_blood_=0.5*·*10^−4^ m s^−1^ (bottom) and for *P* =0.1*·*10^−7^ m s^−1^ (left) and *P* =100*·*10^−7^ m s^−1^ (right) in the presence of active influx, i.e. for various values of *T*_m-in_ (*T*_m-out_=0). Note that the vertical scale is the same in all plots. Fig 7 shows how *P* and *v*_blood_ affect the impact of *T*_m-in_ on brain ECF PK. A smaller value of *v*_blood_ only slightly reduces *C*_ECF_ when *T*_m-in_ is sufficiently high (*T*_m-in_*≥*10·10^−7^ *µ*mol s^−1^), see Fig 7, left. An increase in *P* does reduce the impact of *T*_m-in_ on *C*_ECF_ substantially (Fig 7, right). When the BBB is very permeable, active influx needs to be fast to have any effect, as drug can easily pass the BBB to flow back into the blood plasma. As shown in Fig 7, right, in the presence of a high value of *P, T*_m-in_ only (slightly) affects *C*_ECF_ when it is 10·10^−7^ *µ*mol s^−1^ or higher.

**Fig 7.**
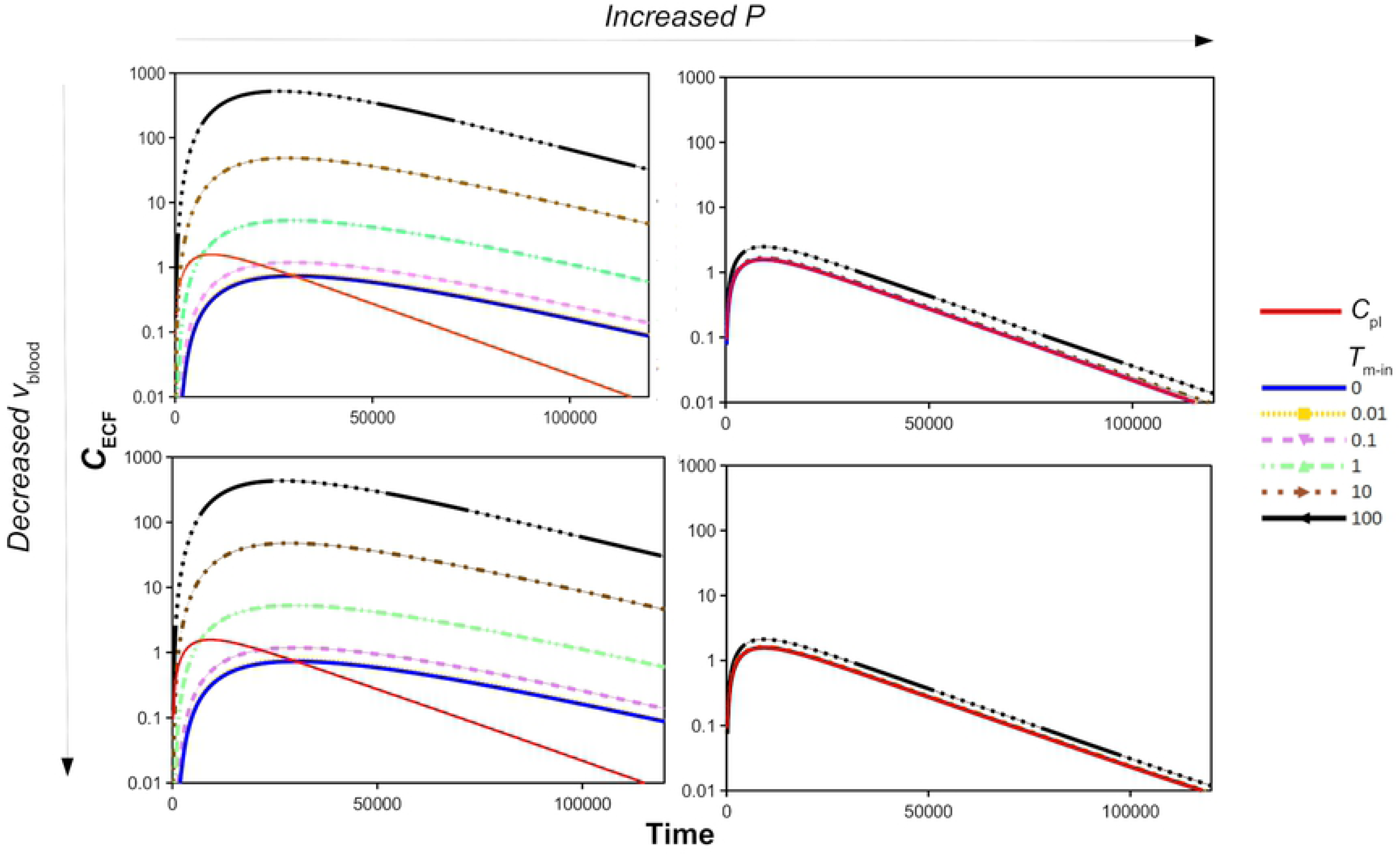
The log concentration-time profiles of unbound drug in brain ECF (*C*_ECF_) with 1000x increased permeability *P* (left to right, 0.1*·*10^−7^ m s^−1^ to 100*·*10^−7^ m s^−1^) or 10x decreased flow *v* _ECF_ (top to bottom, 5·10^−4^m s^−1^ to 0.5·10^−4^ m s^−1^) in the presence of active influx compared to the concentration of unbound drug in the blood plasma (*C*_pl_, red curve). The value of of *T*_m-in_ is changed from 0 to 100·10^−7^ *µ*mol s^−1^, as depicted by various colours. The rest of the parameters are as in Table 2.

Fig 8 shows the log profiles of *C*_ECF_ for *v*_blood_=5·10^−4^ m s^−1^ (top) and *v*_blood_=0.5·10^−4^ m s^−1^ (bottom) and for *P* =0.1·10^−7^ m s^−1^ (left) and *P* =100·10^−7^ m s^−1^ (right) in the presence of active efflux, i.e. for various values of *T*_m-out_ (*T*_m-in_ =0). Fig 8 reveals that *v*_blood_ does not affect the impact of *T*_m-out_ on *C*_ECF_. This is expected, as *v*_blood_ mainly affects *C*_pl_, while active efflux depends on *C*_ECF_. The passive permeability *P* does affect the impact of *T*_m-out_ on *C*_ECF_. If *P* is high, drug can easily flow across the BBB back into the brain ECF, following the concentration gradient between the blood plasma and the brain ECF, thereby countering the effect of *T*_m-out_. Fig 8 (top right) shows that for a high *P, C*_ECF_ is only affected by *T*_m-out_ when its value is higher than 10*·*10^−7^ *µ*mol s^−1^. The values of *C*_ECF_ in the presence of active efflux and a high passive BBB permeability, *P*, are unaffected by *v*_blood_ (Fig 8, right).

**Fig 8.**
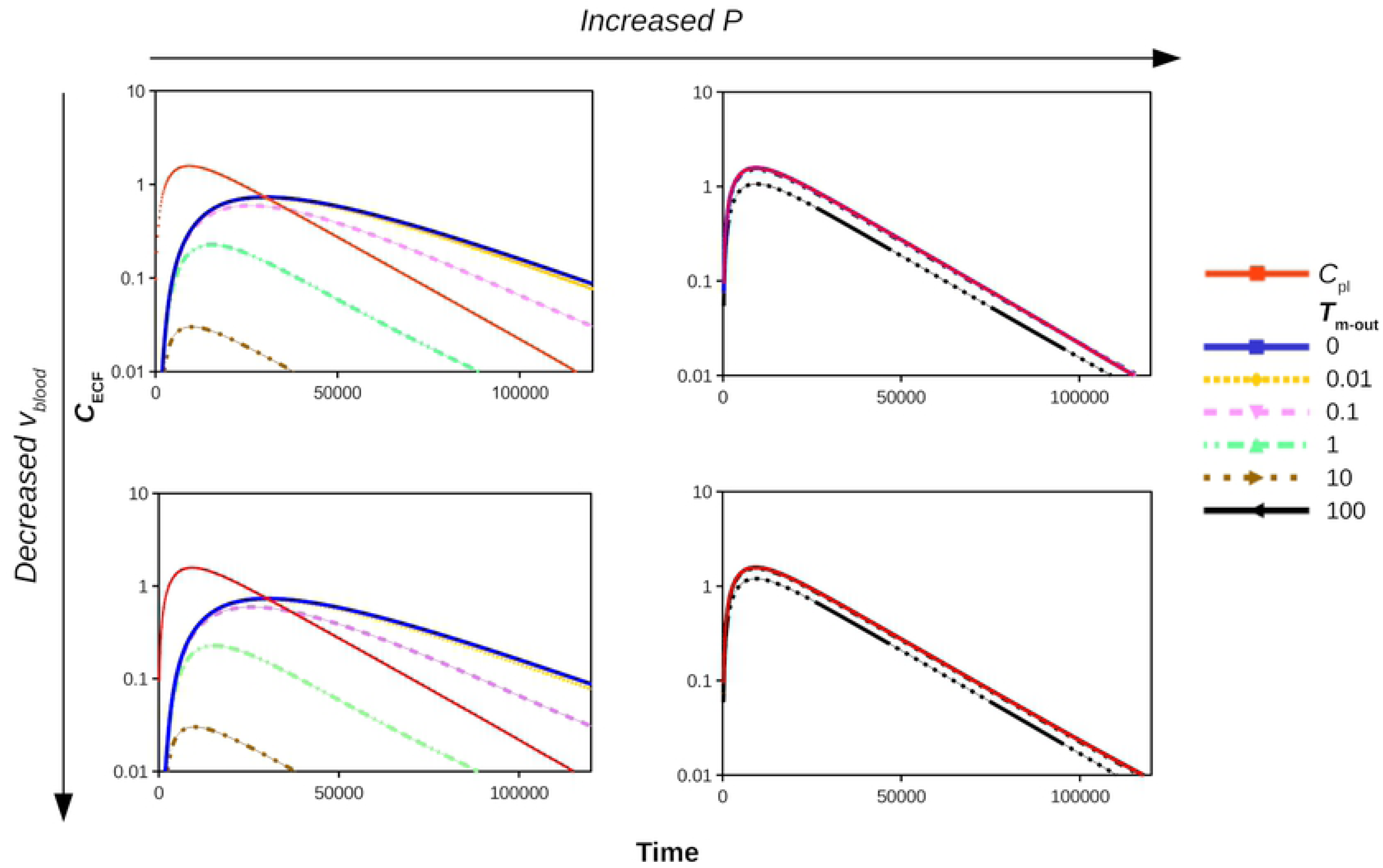
The PK on log-scale of unbound drug in brain ECF (*C*_ECF_) with 1000x increased permeability *P* (left to right, 0.1*·*10^−7^ m s^−1^ to 100*·*10^−7^ m s^−1^) and 10× decreased blood flow velocity *v*_blood_ (top to bottom, 5·10^−7^ m s^−1^ to 0.5·10^−7^ m s^−1^) in the presence of active efflux compared to the concentration of unbound drug in the blood plasma (*C*_pl_, red curve). The value of *T*_m-out_ is changed from 0 to 100*·*10^−7^ *µ*mol s^−1^, as indicated by the different colours. The rest of the parameters are as in Table 2.

Next, we study how the drug distribution within the 3D brain unit is affected by *v*_blood_, *P, T*_m-in_ and *T*_m-out_. Fig 9 shows cross-sections (for 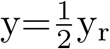 and z=0) of the 3D brain unit at t=5, in which the distribution of *C*_pl_ and *C*_ECF_ is plotted. The values of *C*_pl_ and *C*_ECF_ are represented by shades of red and blue, respectively, where darker shades indicate higher concentrations. In Fig 9a (left) we give a plot for a default *P* and *v*_blood_ (Fig 9a, left). Then, we decrease *v*_blood_ (Fig 9a, middle) or increase *P* (Fig 9a, right). For a lower *v*_blood_, relative differences of *C*_pl_ over space increase (Fig 9a, middle). Additionally, due to the decrease in *C*_pl_, local differences in *C*_ECF_ become more apparent. A larger value of *P* results in an increased exchange of drug between the blood plasma and the brain ECF, such that *C*_ECF_ becomes higher (Fig 9a, right). Fig 9b shows that the presence of active influx (*T*_m-in_=1·10^−7^ *µ*mol s^−1^) increases *C*_ECF_. As a consequence, local differences within *U*_ECF_ become relatively small. With a low value of *v*_blood_, local differences in *U*_pl_ become apparent (Fig 9b, middle). Finally, Fig 9c shows that with active efflux, *C*_ECF_ becomes smaller than when no active efflux is present, except for when *P* is high and more pronounced.

**Fig 9.**
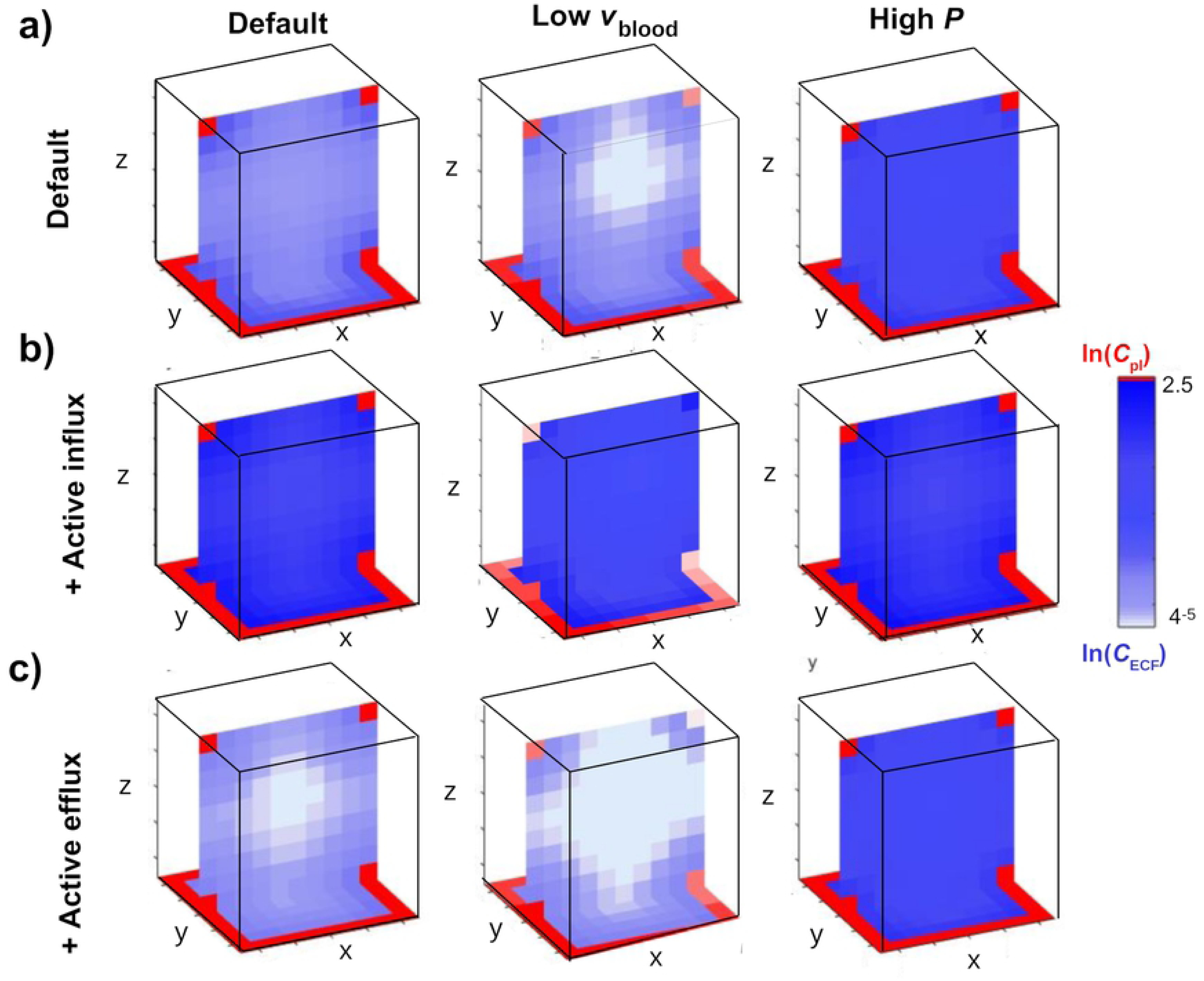
The distribution profiles at cross-sections (at 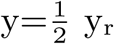) of the 3D brain unit at t=5 of unbound drug in brain ECF with lower brain capillary blood flow velocity (*v*_blood_=0.5·10^−4^ m s^−1^, middle column), higher passive BBB permeability (*P* =100·10^−7^ m s^−1^, right column), presence of active influx (middle row, *T*_m-in_=1·10^−7^ *µ*mol s^−1^) and presence of active efflux (bottom row, *T*_m-out_=1·10^−7^ *µ*mol s^−1^) at t=5. Parameters are as in Table 2.

Values of *C*_ECF_ are given in the table in Fig 10c in order to show the differences within the 3D brain unit more clearly. There, values of *C*_ECF_ are given for four different locations within the 3D brain unit for several values of *v*_blood_ and *P* and t=500. The table again (as in Fig 7, 8 and 9) shows that *v*_blood_ and *P* affect the impact of *T*_m-in_ and *T*_m-out_ on *C*_ECF_. It provides additional information on the distribution of *C*_ECF_ within the 3D brain unit. In general, *C*_ECF_ is higher in the corners relative to the edge and middle within the 3D brain unit. The extent of these local concentration differences depends on the values of *T*_m-in_ and *T*_m-out_. The differences are largest when *T*_m-out_=1 10^−7^ *µ*mol s^−1^, depicted in the lowest line of each sub-table. There, *C*_ECF_ in corner 2 is higher than in corner 1. In addition, in the presence of active influx, the values of *C*_ECF_ are lower in corner 2 than in corner 1. Again, the extent of this difference depends on the value of *T*_m-in_.

**Fig 10.**
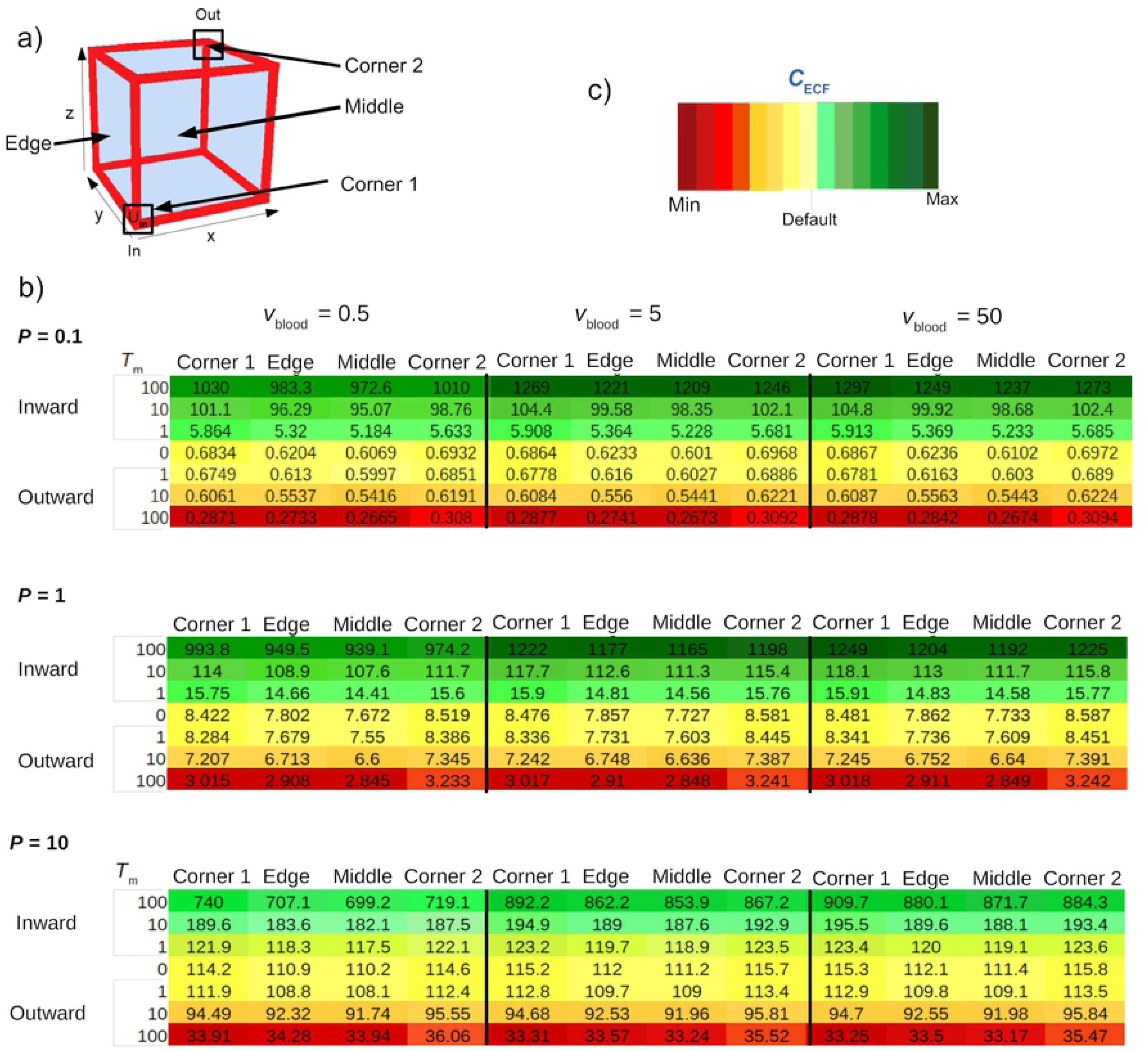
Values of *C*_ECF_ (10^**-3**^ *µ*mol L^−1^) at several locations within the brain unit for different values of *P* and *v*_blood_ at t=500. a) Locations within the 3D brain unit. Corner 1: (x,y,z)=(r,r,r), Corner 2: (x,y,z)=(xr-r,yr-r,zr-r), Edge: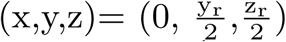, Middle: 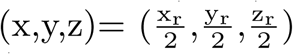. b) Values of *C*_ECF_ are shown for a low ((*P* =0.01·10^−8^m s^−1^), default (*P* =0.1·10^−8^ m s^−1^) and high (*P* =1·10^−8^ m s^−1^) value of *P* in the top, middle and bottom table, respectively. Within each table, concentrations are given for several values of *v*_blood_ (*v*_blood_=0.5·10^−4^ m s^−1^, *v*_blood_=5·10^−4^ m s^−1^ and *v*_blood_=50·10^−4^ m s^−1^, left to right), *T*_m-in_ (*T*_m-in_=0, *T*_m-in_=1·10^−7^ *µ*mol s^−1^, *T*_m-in_=10·10^−7^ *µ*mol s^−1^ and *T*_m-in_=100·10^−7^ *µ*mol s^−1^) and *T*_m-out_ (*T*_m-out_=0, *T*_m-out_=1·10^−7^ *µ*mol s^−1^, *T*_m-out_=10·10^−7^ *µ*mol s^−1^ and *T*m-out=100·10^−7^ *µ*mol s^−1^) at different locations. When *T*_m-in_ is changed, *T*_m-out_=0 and vice versa. c) Colour legend. In each table, colours are relative to the value of *C*_ECF_ in the middle of the unit in the absence of active transport for *v*_blood_=5*·*10^−4^ m s^−1^, of which the colour is denoted by “Default”. The intensity of green corresponds to the extent of increase, and the intensity of red corresponds to the extent of decrease of *C*_ECF_ compared to the default. Other parameters are as in Table 2.

## 4 Discussion

We have developed a mathematical model that describes the local distribution of a drug within a 3D brain unit as an extension of our earlier 2D proof-of-concept model [29]. The 3D brain unit is represented as a cube. This new model provides an important step towards more realistic features of the brain. The 3D representation allows for the representation of the brain ECF as continuous. The brain capillary blood flow and active transport across the BBB have been explicitly incorporated. This enables us to more realistically predict the impact of the interplay of cerebral blood flow, BBB characteristics, brain ECF diffusion, brain ECF bulk flow and brain (target) binding on drug distribution within the brain. Altogether our model allows the study of the effect of a large amount of parameters values (summarized in Table 1) on drug distribution within the 3D brain unit.

This study has focused on the effect of the newly implemented brain properties on brain ECF concentrations a drug within the brain. It is shown that the brain capillary blood flow velocity and the passive BBB permeability affect the concentration of a drug within the brain, and, as anticipated [68, 69] that a low brain capillary blood flow velocity affects the short-term, but not the long-term concentration-time profiles of *C*_pl_ and *C*_ECF_, (Fig 3 and 4). Also, passive BBB permeability has a high impact on brain ECF PK, even when drug is actively transported across the BBB. Moreover, the BBB permeability and, in smaller extent, the brain capillary blood flow velocity affect the impact of active influx on drug PK within the brain ECF (Fig 7 and 8). Interestingly, the brain capillary blood flow velocity, passive BBB permeability and active transport do not only affect the concentration of drug within the brain ECF, but also its distribution within the brain ECF (Fig 9 and 10.

Taken together, the 3D brain unit model shows the impact of drug-specific and brain-specific parameters on drug distribution within the brain ECF. The added value is that all these factors can now be studied *in conjunction* to understand the interdependencies of multiple brain parameter values and drug properties. This makes this single 3D brain unit model suitable for the next step, which is to mount up multiple units to represent a larger volume of brain tissue, in which the brain tissue properties for each unit can be defined independently. The units may be given different systemic properties (such as the BBB permeability or drug target concentration), to represent the heterogeneity of the brain in a 3D manner.

## S1 Appendix Nondimensionalization of the model

We can make Eq (2–16) dimensionless by introducing a change of variables. Here, the original variables are scaled to dimensionless variables by scaling with a characteristic, dimensional scale. We set:

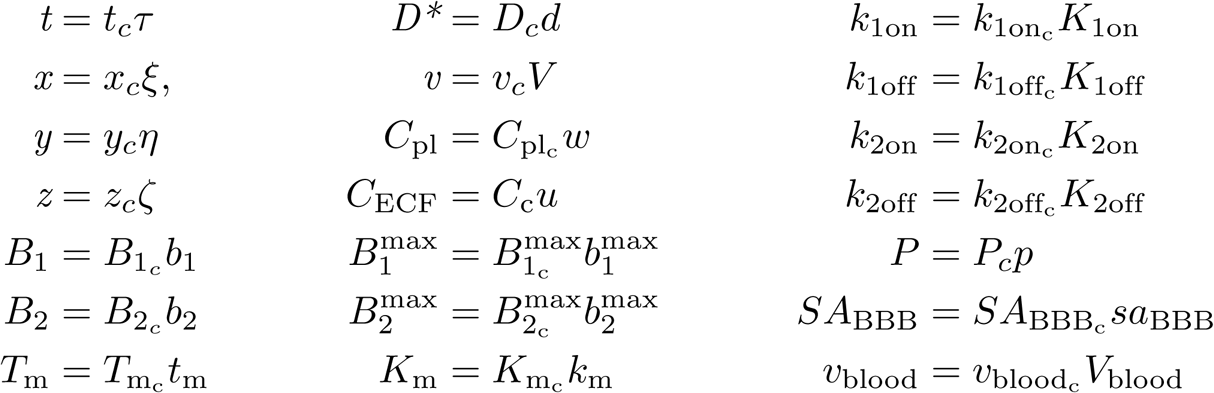

where

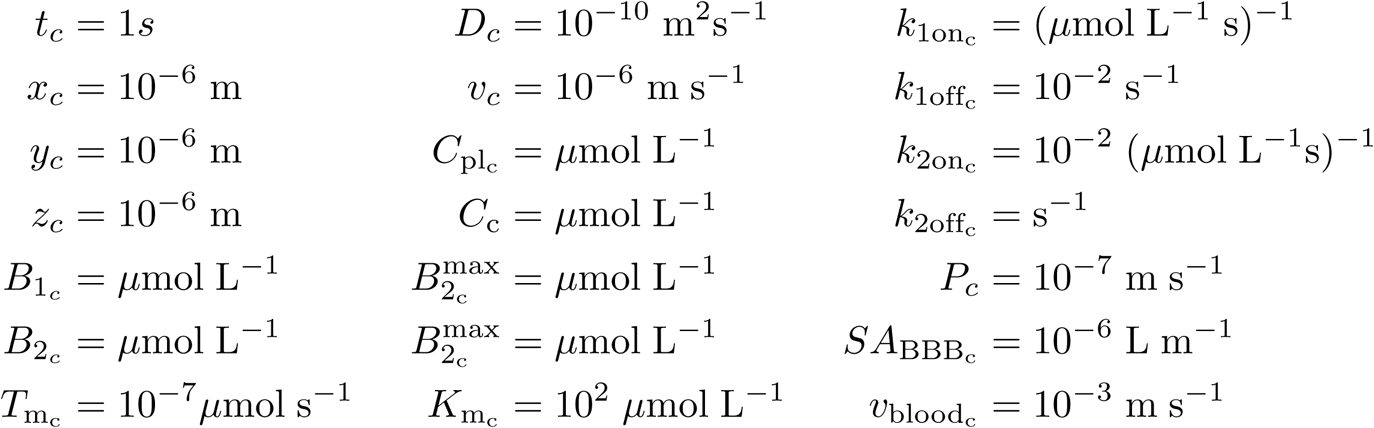

This leads to the following dimensionless equation for drug in the blood plasma (example based on Eq (2), but similar for Eq (3)–(4)):

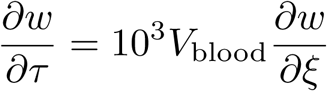

and the following system of dimensionless equations for drug within the brain ECF (for Eq (6)):

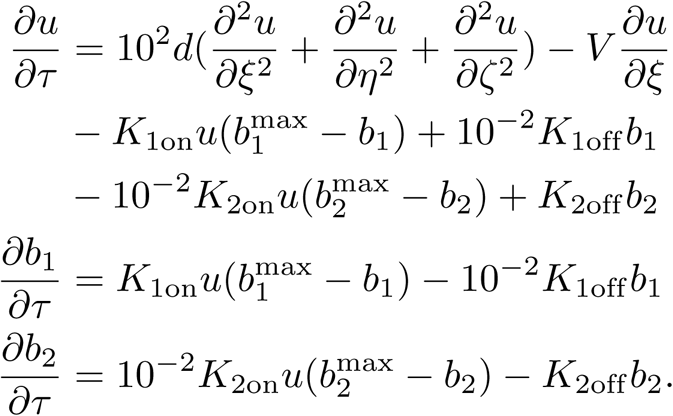

The corresponding boundary conditions (Eq (10)–(11), example for Eq (10), but similar for Eq (10)) are given by:

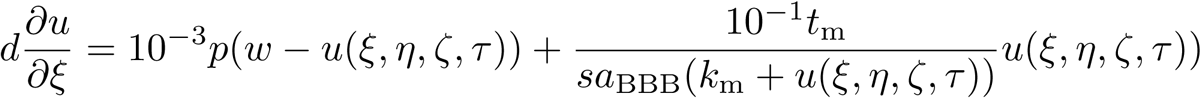

for ξ=0 and ξ=1.

The initial conditions become

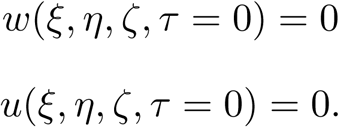

## S2 Appendix The effect of paracellular permeability on PK within the brain ECF

We study the passive transcellular permeability and passive paracellular permeability separately. This is different from before, where we have studied the total passive permeability. We study the effect of paracellular transport on the PK within the 3D brain unit. The paracellular permeability can increase due to disruption of the BBB, which in turn could be a result of disease. We include paracellular permeability and study its effect on drug concentrations within the 3D brain unit. For drugs for which the passive transcellular BBB permeability is low (*P*_trans_=0.01*·*10^−7^ *µ*mol s^−1^), increasing has a large impact on drug PK within the brain (Fig 1, left). For drugs with a high passive permeability, an increased paracellular permeability has less effect, as shown in Fig 1(right). Essentially, changing the paracellular permeability has a similar effect as changing the total and the transcellular permeability: both increase the transport of drug along the concentration gradient between the blood plasma in the brain capillaries and the brain ECF.

**Supplementary Figure 1.**
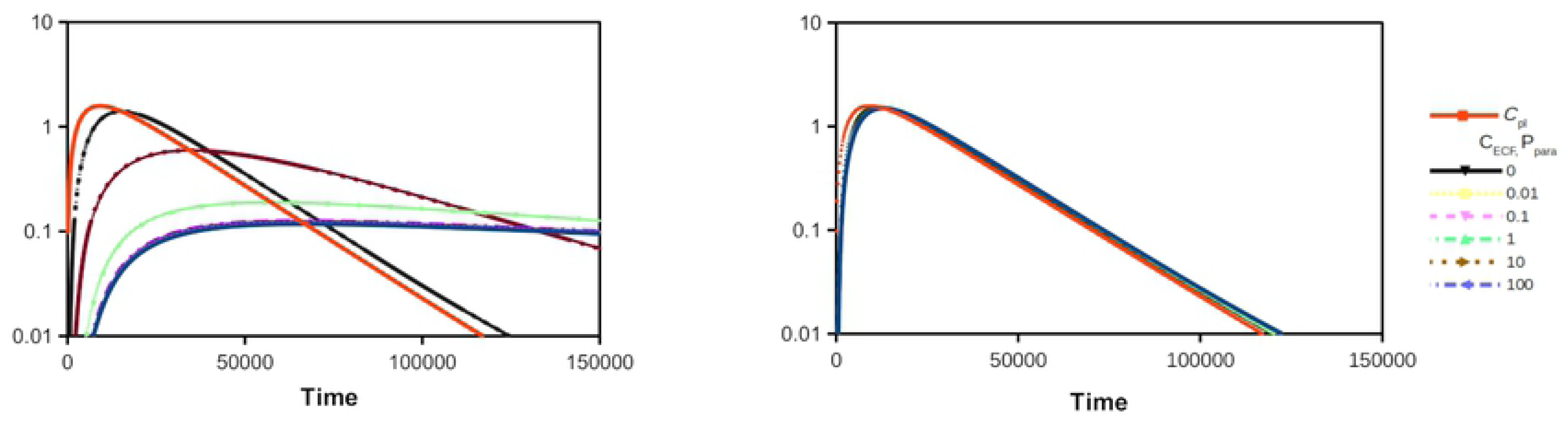
The PK in log-scale of unbound drug in the brain ECF (*C*_ECF_) compared to the concentration of unbound drug in the blood plasma (*C*_pl_, red curve). The transcellular passive permeability, *P*_trans_, is set to 0.01·10^−7^ m s^−1^ (left) and 1·10^−7^ m s^−1^ (right), while the paracellular permeability, *P*_para_ is changed from 0 to 1·10^−1^ m s^−1^ as depicted by different colours.

## S3 Appendix The Renkin-Crone equation and the 3D brain unit model

We compare our model with the Renkin-Crone equation, which is a well-known equation relating blood flow to tissue uptake [64, 65], see Box I. The Renkin-Crone equation predicts that the transport of drugs across the BBB *into* the brain depends on the brain capillary blood flow rate, *Q*, in the presence of a large BBB permeability surface, *PS*. The volumetric parameters *Q* and *PS* are related to the brain capillary blood flow velocity, *v*_blood_, and the BBB permeability, *P*, by the brain capillary and BBB surface area, *SA*_cap_ and *SA*_BBB_, respectively. Here, we study the effect of *v*_blood_ on the passive transport of drug into the brain for different values of *P*. For this purpose, we:

1. Take a constant concentration of drug within the blood-plasma-domain (i.e. we set *C*_pl_(t)=1 for *C*_pl_(t)∈ *U*_in_.
2. We simplify boundary conditions (11) and (12) to 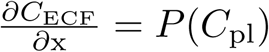 in order to study passive *influx*, which is the passive movement of drug *into* the brain, only. Note that his is different from the approach we took previously, in which passive transport into or out of the brain ECF depends on a difference in concentration between the blood plasma and the brain ECF (see Eq(10)). Moreover, we set *T*_m-in_=0 and *T*_m-out_=0
3. We leave out drug binding and set *B*_1_^max^, *B*_2_^max^=0.

#### Box I The Renkin-Crone equation

The brain capillary blood flow affects the passive clearance of a drug across the BBB according to the Renkin-Crone equation [64, 65]. The Renkin-Crone equation describes the relation between the brain capillary blood flow and transport across the BBB as follows:

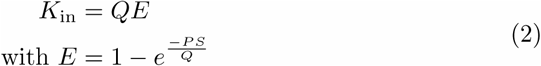

with *K*_in_ the passive clearance of drug from the blood into the brain (L s^−1^), *Q* (L s^−1^) the blood flow rate in the brain capillaries and *PS* (L s^−1^) the passive permeability surface of the BBB. Both *Q* and *PS* have the same units, such that, *E*, the ratio of compound extracted from the blood into the brain, is dimensionless. The Renkin-Crone equation shows that the transport from the blood into the brain depends on the ratio of the BBB permeability surface (*PS*) and the blood flow rate (*Q*). When *PS* ≫*Q*, the extraction ratio *E* approaches 1, such that *K*_in_ is determined by changes in *Q*. In other words, when *PS* ≫*Q*, drug transport across the BBB is much faster than the rate of drug supply into the brain capillaries. Then, drug transport into the brain can only be increased by increasing *Q*. On the other hand, when *Q* ≫*PS, E* approaches 0. In this case, the drug supply into the brain capillaries is much faster than the rate of drug transport across the BBB. Then, drug transport into the brain can only be increased by increasing *PS*. The Renkin-Crone equation implies that the effect of the brain capillary blood flow rate on the concentration of unbound drug exchanging with the brain is most pronounced for drugs that easily cross the BBB [65, 70], i.e. drugs for which *PS* ≫*Q*, or, in terms of velocity rather than rate, drugs for which *P* ≫*v*_blood_. Under general, non-pathological circumstances *v*_blood_ is around 5·10^−4^ m s^−1^ (see Tables 1 and 2), which implies that BBB transport is impacted by the blood flow velocity when drug molecules have a value of *P* that is (much) higher than 10^−4^ m s^−1^ (i.e. 10^3^ times the default value as given in Table 2).

We measure the change in 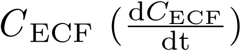 at one specific point of the 3D brain unit, 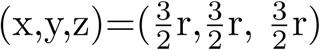,which we denote by *u*_1_, as indicated in Fig 2 (top). Similarly, we measure *C*_pl_ at one specific point of the 3D brain unit, 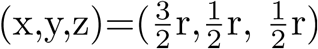, denoted by *w*_1_, as indicated in Fig 2 (top). It takes some time until a steady state is reached and values of *C*_pl_ and 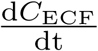 are approximately constant, see Fig 2 (bottom). At steady state *k*_BBB_, which is the rate constant of drug transport from the blood plasma across the BBB into the brain ECF, can be determined as follows:

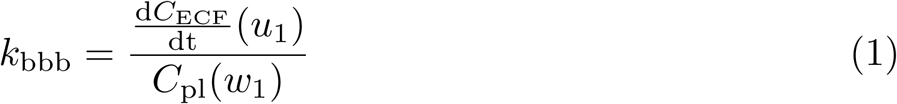

with 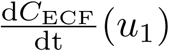 the change in *C*_ECF_ over time in *u*_1_ and *C*_pl_(*w*_1_) the value of *C*_pl_ in *w*_1_ when both *C*_pl_(*w*_1_) and 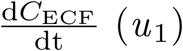 do not longer vary. Fig 3 demonstrates that the way *v*_blood_ affects *k*_BBB_ varies with the value of *P*. With values of *P* of 1*·*10^−4^ m s^−1^ or lower, *k*_BBB_ is independent of *v*_blood_. With values of *P* of 10·10^−4^ m s^−1^ or higher, *k*_BBB_ linearly increases with *v*_blood_ up to a certain threshold (e.g. for *P* =10·10^−4^ m s^−1^, *k*_BBB_ starts to approach constant levels when *v*_blood_ ≥2). These results correspond to the predictions of the Renkin-Crone equation (Box I).

**Supplementary Figure 2.**
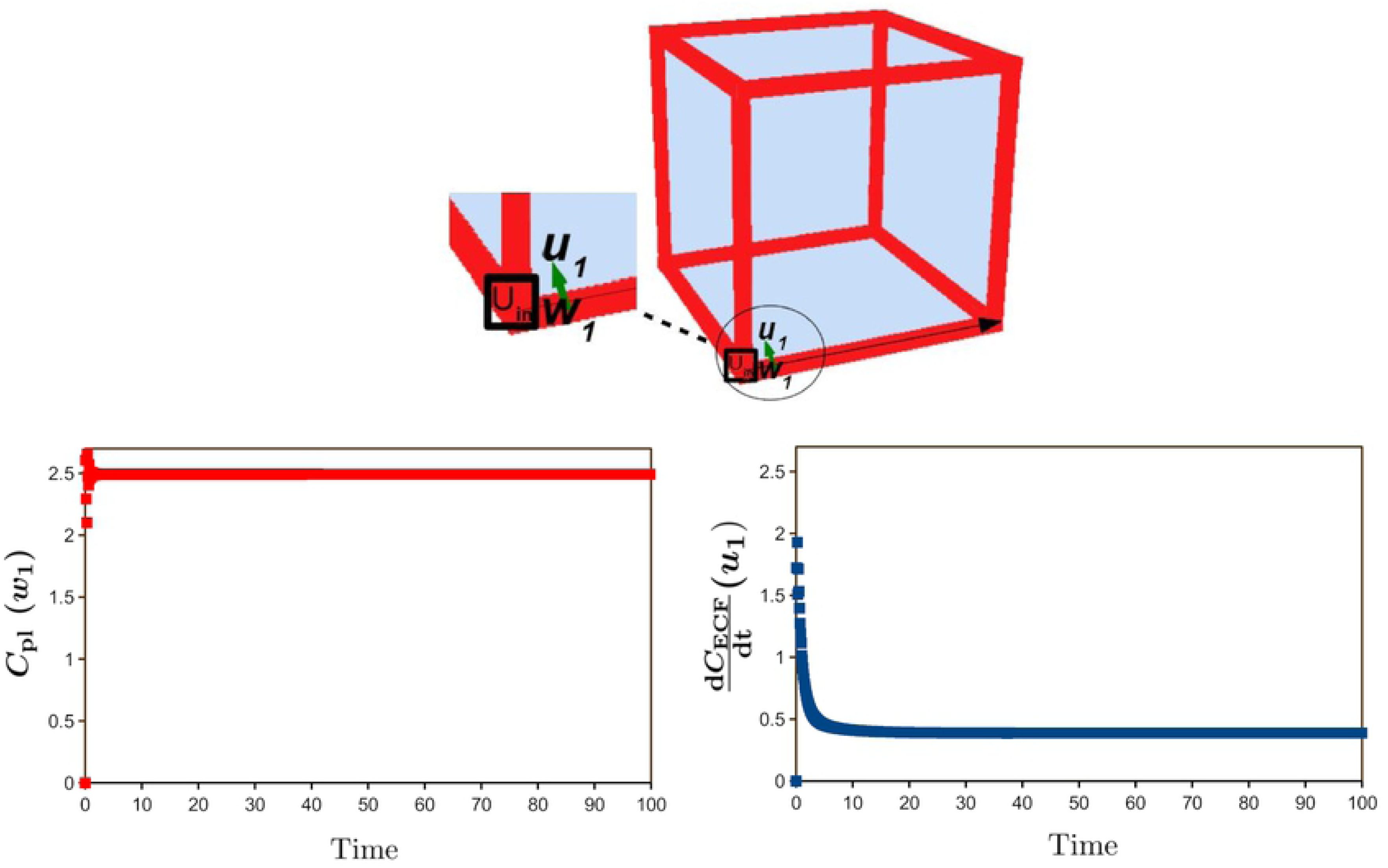
Determination of *C*_pl_(*w*_1_) and 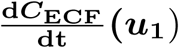. Time Top: Locations of *w*_1_ and *u*_1_, where *C*_pl_(*w*_1_)and 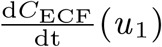 are measured, within the 3D brain unit. The black arrow indicates the direction of the brain capillary blood flow, while the green arrow indicates the direction of BBB transport. Bottom: Profiles of *C*_pl_ (*w*_1_) and 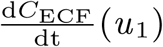 over time.

**Supplementary Figure 3.**
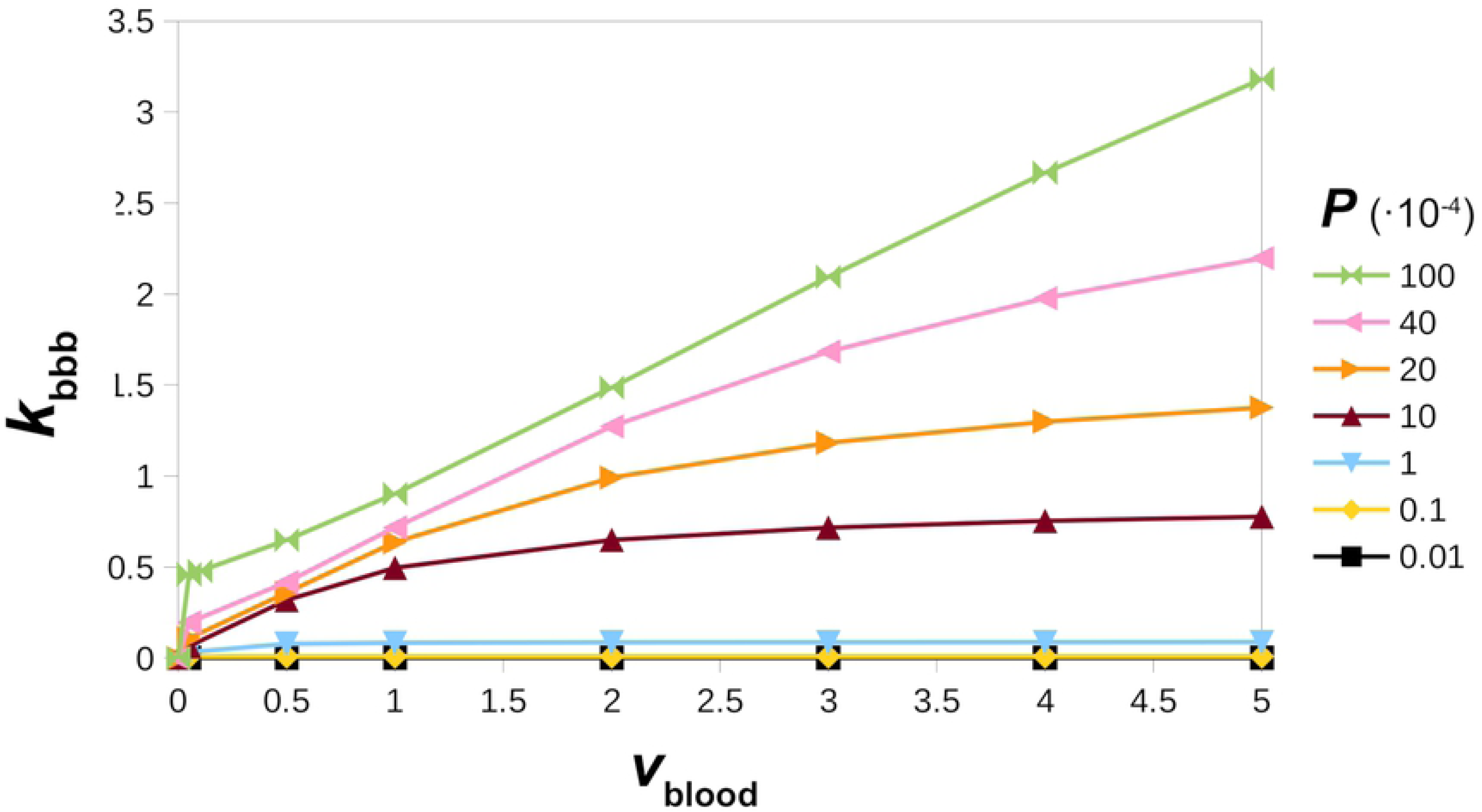
The effects of *v*_blood_ on *k*_BBB_. The effect of *v*_blood_ on *k* BBB depends on *P*. Note that here *P* is taken 10^3^ times its default value, see Table 2.

